# Erythrocyte Count, Anemia, and the Human Natural Lifespan Limit: Evidence from the Long Life Family Study

**DOI:** 10.64898/2026.06.08.730977

**Authors:** Konstantin G. Arbeev, Olivia Bagley, Joanne M. Murabito, Harvey J. Cohen, Brandon L. Pierce, W. Evan Johnson, Dan T.A. Eisenberg, Milind C. Mahajan, Stacy L. Andersen, Kaare Christensen, Joseph M. Zmuda, Bharat Thyagarajan, Sheng Luo, Anatoliy I. Yashin, Michael A. Province, Abraham Aviv

## Abstract

Erythropoiesis is the most replication-intensive process in the body. Its lifelong replicative demands may erode hematopoietic cells’ replicative capacity, leading to a decline in erythrocyte count (EC) in older adults and limiting their lifespan. We examined the relationship between EC and mortality among 1,620 participants aged ≥70 years in the Long Life Family Study and found that lower EC further augmented the exponential, age-dependent rise in mortality. We identified an EC threshold (ECT) (3.8x10^12^/L) below which mortality was amplified (p=9.3x10^-6^). As EC declined with age (p=8.2x10^-18^), it fell below this threshold in many participants, sharply increasing their mortality risk. This mortality-based ECT emerged from modeling, independent of the World Health Organization anemia definition, which is based on statistical thresholds (5^th^ centiles) of hemoglobin distribution in populations ≤ 65 years. Thus, declining EC may be one of the biological factors imposing a natural lifespan limit on many older adults.

---

Life expectancy rose by nearly 30 years during the 20th century, but the rate of increase has recently decelerated in the longest-living populations ^1^. One possible explanation is a biological limit on the replicative capacity of human somatic cells. This was discovered by Hayflick and Moorhead ^2^, who found that cultured human somatic cells replicate only a finite number of times before entering senescence, a state of permanent growth arrest known as the Hayflick limit ^3^.

Erythropoiesis, the most replication-intensive process in the human body, is the central driver of hematopoiesis ^4^. Should the Hayflick limit function *in vivo*, the replicative capacity of hematopoietic stem cells might diminish with age ^5^. The resulting impairment of erythropoiesis could contribute to anemia of aging and its associated increase in mortality risk ^6–9^.

Hemoglobin (Hgb) concentration below the 5^th^ centiles of the Hgb distribution in reference populations ≤ 65 years is defined as anemia by the World Health Organization (WHO) ^10,11^. Such a dichotomous, distribution-based statistical definition does not capture the age-related decline in erythrocyte count (EC) and its potential outcomes as older adults approach a natural lifespan (NLS) limit, a putative maximum age beyond which survival is biologically constrained. In this study, we have examined relationships between EC and a well-defined outcome, all-cause mortality (hereafter, mortality), in the National Institute on Aging Long Life Family Study (LLFS) ^12^. By design, the LLFS is enriched for families with a higher probability of exceptional longevity, and thus with a greater likelihood than the general population of nearing the NLS limit.

## RESULTS

We applied a Cox proportional hazards model (CPHM), a joint model (JM), and a stochastic process model (SPM) (**Methods**) to: (a) assess whether EC modifies the age-related increase in mortality risk among LLFS participants aged 70 years or older (using a single measure of EC per participant); (b) examine the associations between participant-specific EC trajectories, defined by baseline EC and its age-related change, and mortality; (c) test for the presence of an EC threshold (ECT) below which mortality risk increases; and (d) determine sex effects on EC trajectories and their associations with mortality and survival.

### Erythrocyte count trajectories past the age of 70 years

The sample comprised 1,620 LLFS participants without cancer or kidney disease, with one or more EC measurements at age 70 years or older, and with information on mortality and key covariates (age and sex) used in the main models (Methods). Table 1 summarizes participants’ characteristics. Extended Data Figure 1 displays participant selection. Supplementary Table S1 provides information on additional covariates (smoking, body mass index, cystatin C, high-sensitivity C-reactive protein, interleukin-6, and transferrin receptor) for the entire sample and for a subsample of 520 participants with two or three sequential EC measurements.

**Table 1.** Characteristics of the Long Life Family Study participants in this study.

| **Characteristics** | **Females** | **Males** | **Total Sample** |
| --- | --- | --- | --- |
| Number of participants | 937 | 683 | 1,620 |
| Number of participants with 2 or 3 observations | 298 | 222 | 520 |
| Number (%) of deceased participants | 437 (46.6%) | 328 (48.0%) | 765 (47.2%) |
| Follow-up period (years) (mean ± SD) | 10.4 ± 5.2 | 10.2 ± 5.4 | 10.3 ± 5.3 |
| Age at first observation (mean ± SD) | 82.4 ± 9.5 | 81.5 ± 8.9 | 82.0 ± 9.2 |
| US participants (%) | 67.7% | 65.2% | 66.6% |
| EC at ages 70 years and older<sup>a</sup> | 4.40 ± 0.45 | 4.60 ± 0.56 | 4.48 ± 0.51 |
| EC at ages 70-79 years<sup>a</sup> | 4.51 ± 0.41 | 4.80 ± 0.47 | 4.64 ± 0.46 |
| EC at ages 80-89 years<sup>a</sup> | 4.36 ± 0.38 | 4.57 ± 0.51 | 4.45 ± 0.45 |
| EC at ages 90-99 years<sup>a</sup> | 4.25 ± 0.48 | 4.27 ± 0.57 | 4.26 ± 0.52 |
| EC at ages 100 years and older<sup>a</sup> | 4.09 ± 0.59 | 3.81 ± 0.52 | 4.02 ± 0.58 |
| Annual rate of change in EC at ages 70 years and older<sup>b</sup> | -0.016 ± 0.054 | -0.025 ± 0.059 | -0.020 ± 0.056 |
| Annual rate of change in EC at ages 70-79 years<sup>b</sup> | -0.011 ± 0.054 | -0.012 ± 0.051 | -0.011 ± 0.052 |
| Annual rate of change in EC at ages 80-89 years<sup>b</sup> | -0.019 ± 0.064 | -0.030 ± 0.076 | -0.024 ± 0.069 |
| Annual rate of change in EC at ages 90 years and older<sup>b</sup> | -0.023 ± 0.062 | -0.040 ± 0.090 | -0.031 ± 0.076 |

<sup>a</sup> mean ± SD, $10^{12}/L$ ; <sup>b</sup> mean ± SD, $10^{12}/L$ per year. EC: erythrocyte count. The annual rate of change in EC is computed for 520 participants with at least 2 EC measurements. For participants with two visits, $\Delta EC$ was computed as $\Delta EC = (EC_{visit2} - EC_{visit1}) / (Age_{visit2} - Age_{visit1})$ . For participants with three visits, $\Delta EC$ was computed separately for the intervals between visit 2 and visit 1, and between visit 3 and visit 2.

ECs declined with age in both sexes. On average, females had lower EC than males, but their rate of decline was slower (Figure 1, Table 1). The magnitude of decline was similar between sexes at ages 70–79 years, but was greater in males than in females at ages 80–89 years and 90 years or older (Table 1).

**Figure 1.**
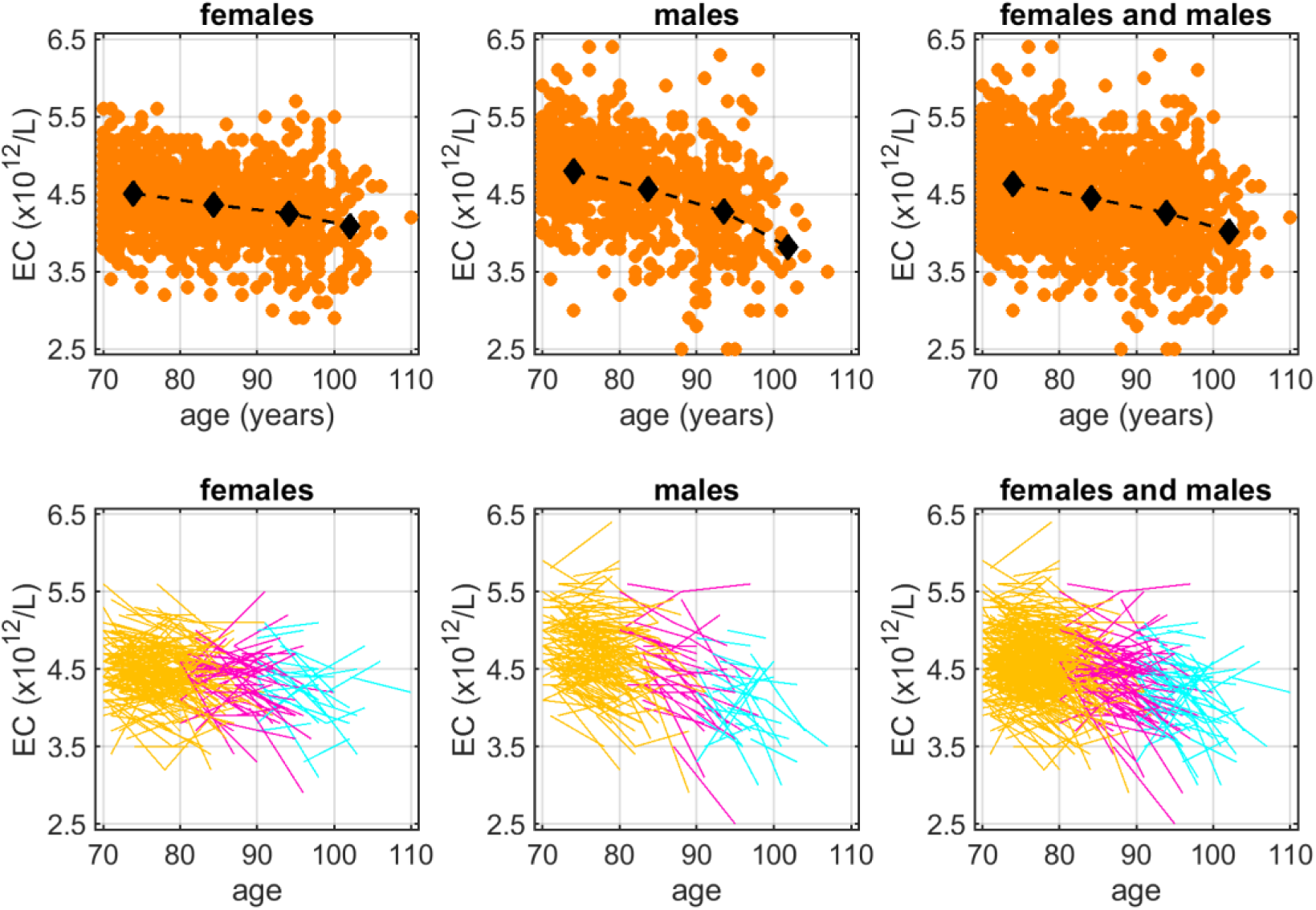
Erythrocyte count in LLFS participants, ages 70 years and older. Top panels: scatter plots of erythrocyte count (EC) in 1,620 participants. Black diamonds denote average EC in different age groups 70-79, 80-89, 90-99, and 100+ years. Bottom panels: individual EC trajectories in a subgroup of 520 participants with two or three sequential ECs. Baseline age groups are color-coded: 70-79 years (orange), 80-89 years (magenta), and 90+ years (cyan). Lines connect baseline and follow-up ECs for the same individuals. Four outliers (three < 2.5 × 10^12^/L and one > 6.5 × 10^12^/L) were removed.

### Association of the first erythrocyte count at age 70 years or older with mortality risk

A total of 765 (47.2%) participants died during the follow-up period (mean ± SD: 10.3 ± 5.3 years; maximum follow-up: 18.4 years; Table 1). Using the CPHM, we found an association between lower EC and increased mortality risk in the main model, adjusting for age and sex. The hazard ratio (HR) per unit decrease in EC (1 × 10^12^/L) was 1.32 (95% confidence interval [CI]: [1.12, 1.60]). This indicates that participants with 1 × 10^12^/L lower EC had a 32% increase in mortality risk compared to those with higher EC, after adjusting for age (HR per one year increase: 1.14, 95% CI: [1.13, 1.15]), and sex (HR for males vs. females: 1.50, 95% CI: [1.29, 1.71]).

Because mortality rises exponentially with age, the association of low EC beyond age 70 years with increased mortality risk was especially striking (Figure 2). The trajectories depict the ratios of mortality risks across different ages (x-axis) and ECs, relative to the mortality risk at age 70 years and the average EC (4.5 × 10¹²/L) at that age. Because both EC and mortality are scaled to age 70 years, the HR for mortality at age 70 is 1 when EC = 4.5 × 10¹²/L, rising to 1.3 when EC = 3.5 × 10¹²/L, and dropping to 0.9 when EC = 5.0 × 10^12^ (Figure 2, inset table). However, at age 90 years, for instance, the HR for mortality is already 13.7 at the average EC, due to the natural increase in mortality with age; it rises further to 18.0 when EC = 3.5 × 10¹²/L and drops to 11.9 when EC = 5.0 × 10^12^.

**Figure 2.**
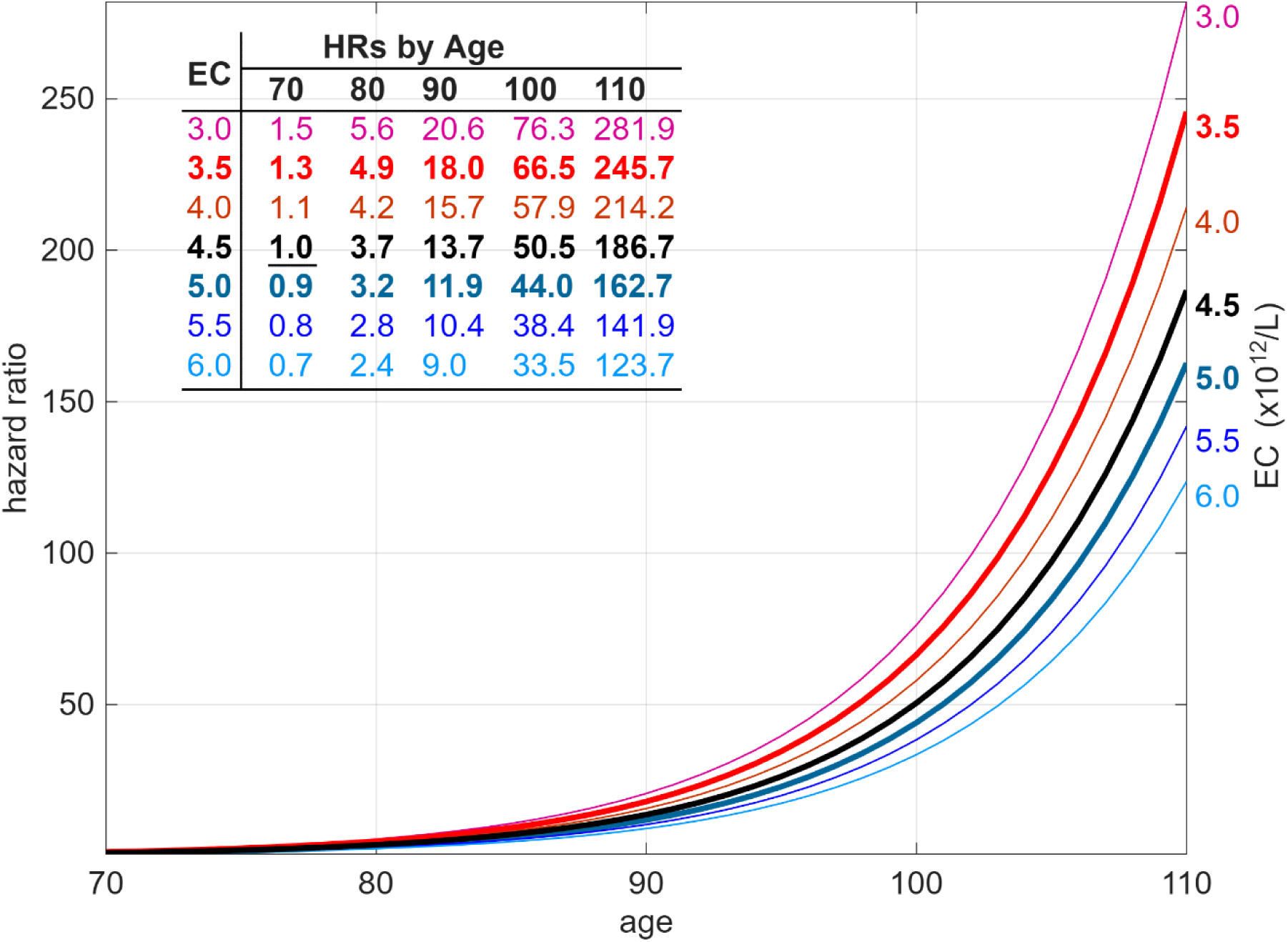
Associations between mortality beyond age 70 years and erythrocyte count (Cox proportional hazards model adjusted for age and sex). Hazard ratios (HRs) for mortality risk are shown across different ages (x-axis) and erythrocyte counts (ECs), relative to the mortality risk at age 70 years and the average EC at that age (4.5 × 10^12^/L). The thick, bold black line shows the HR trajectory for the average EC at age 70 years. Colored lines show HR trajectories for lower EC (above the black line) and higher EC (below the black line). The table presents numerical examples of HRs for specific ages and ECs, scaled to an age of 70 years and an EC of 4.5 × 10^12^/L (<u>underlined</u>). The thick, bold, colored lines correspond to two EC examples (3.5 × 10^12^/L and 5 × 10^12^/L) discussed in the text. The corresponding rows are highlighted (bold) in the inset table.

### Association of erythrocyte count trajectories with mortality risk

We used a JM to examine associations between individual participants’ EC trajectories and time to death (Methods). The regression coefficient for EC trajectories in the hazard function quantifies the association between participant-specific deviations of EC from the expected (population-average) EC and mortality. For this analysis, all available EC measurements from 1,620 participants were used to estimate the joint likelihood function, while within-person EC changes were informed by the 520 participants with repeated EC measurements.

The estimated regression coefficient for EC trajectories was -0.326 (Supplementary Table S2, panel B), indicating that lower subject-specific EC trajectories relative to the model-implied expected trajectory are associated with higher mortality hazard, corresponding to an HR of 1.39 per 1 × 10^12^/L decrease below the expected EC trajectory. Thus, when applied to longitudinal EC trajectories, the JM confirmed the CPHM’s findings for the first EC measurement after age 70 years. The JM also confirmed a declining EC trajectory with age, i.e., a decline of about 0.021 × 10^12^/L per year (“time” in Supplementary Table S2, panel A; Extended Data Figure 2).

### Testing for an erythrocyte count threshold of mortality

The SPM offers more flexible specifications for the relationship between EC and mortality risk than either the JM or the CPHM (Methods and Supplementary Text). Like the JM, the SPM used EC measurement data from all 1,620 participants. The SPM indicated that an EC threshold (ECT) defines the relationship between EC and mortality risk (Methods), such that mortality increases above baseline for the respective age when EC falls below the ECT. For ECs above the ECT, there is no association between EC and mortality, and the risk remains at the baseline level for that age. We modeled the additional mortality risk below the ECT as a quadratic function of EC. We also assumed that the width of this quadratic function may change with age (e.g., narrow) and tested the null hypothesis that the width is independent of age.

The SPM identified an association between low EC and increased mortality (p = 9.3 × 10^-6^), confirming the existence of the ECT. Under the fitted threshold SPM, the estimated ECT was 3.8 × 10^12^/L, with a 95% bootstrap CI of [3.4 × 10^12^/L, 4.1 × 10^12^/L]. Importantly, we found that the quadratic function describing EC-related mortality below the ECT becomes narrower with age (p = 0.007), indicating that the same deviation of EC from the ECT at older age results in a larger increase in mortality risk than at younger ages.

### The sex effect

#### Results based on erythrocyte count at age 70 years or older

The male-female difference in EC (Table 1, Figure 1) may contribute to a sex difference in mortality risk. The CPHM highlights the amplification of mortality risk at lower EC (Figure 2). This amplification translates into different survival probabilities for participants with different starting ECs at age 70 years. Figure 3 shows survival probabilities beyond age 70 years. Notably, the survival probabilities do not differ much for females and males with different ECs at age 75 years and ages beyond 100 years. However, at ages between these extremes, the differences in survival chances between individuals with different starting ECs at age 70 are considerable. For example, according to the model, a female participant with EC = 3.5 × 10¹²/L at age 70 has a 41.5% chance of surviving to age 90, compared with a 60.2% chance for a same-age female participant with EC = 5.5 × 10¹²/L. For a male participant with EC = 3.5 × 10¹²/L at age 70, the chance of surviving to age 90 is 26.8%, whereas it is 46.8% for a same-age male participant with EC = 5.5 × 10¹²/L. The bottom panel of Figure 3 compares survival probabilities for females and males at different ECs and ages.

**Figure 3.**
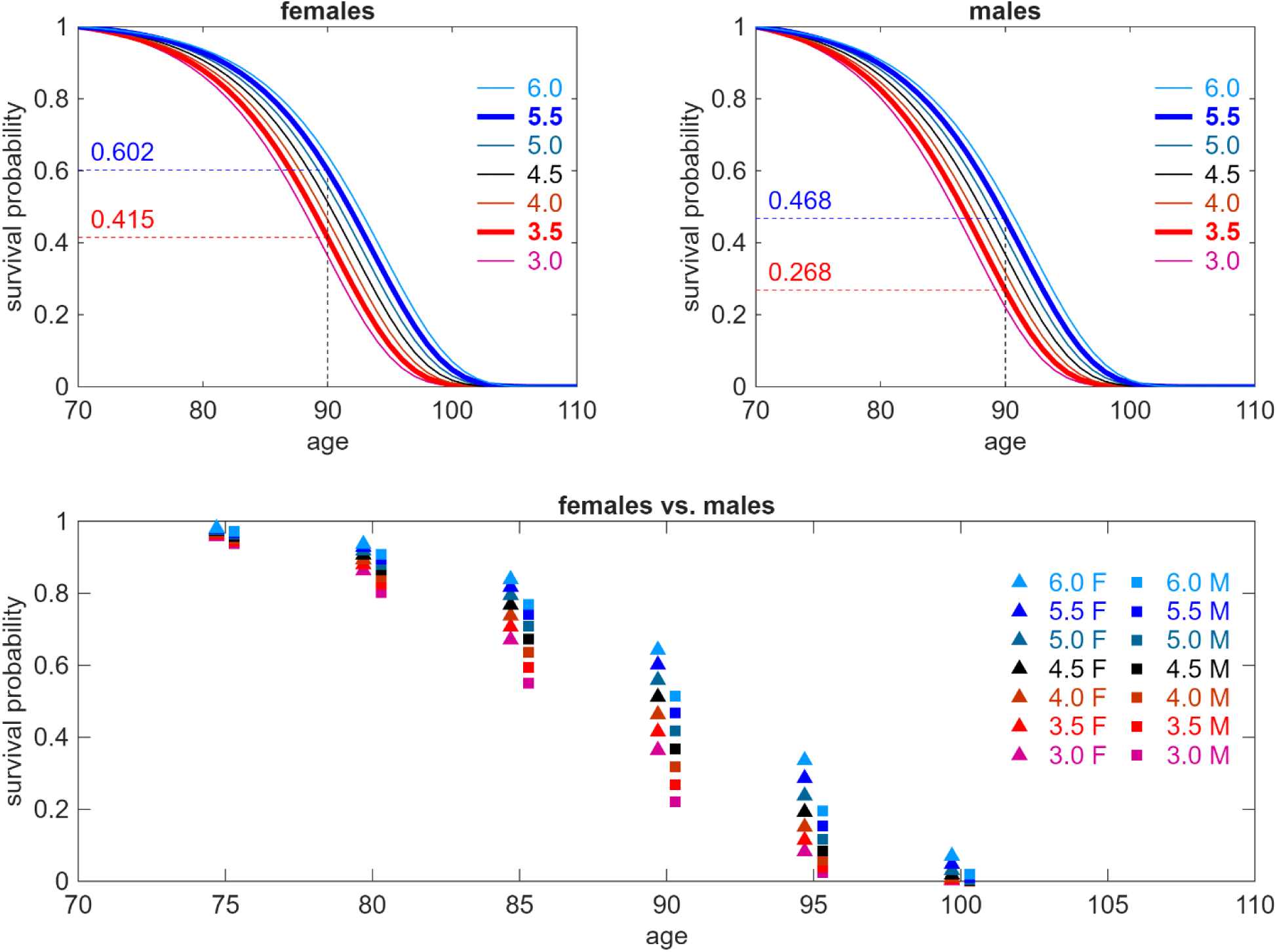
Survival beyond age 70 years in females and males for different erythrocyte counts at age 70 years (estimated from Cox model). Survival functions for females (top left) and males (top right) are shown for different erythrocyte counts (ECs) (x 10^12^/L) at age 70 years. Two thick, bold, colored lines correspond to the EC values of 3.5 × 10^12^/L and 5.5 × 10^12^/L. Examples of survival probabilities are displayed along the y-axis and are color-coded accordingly. The bottom panel compares survival probabilities for females (F) and males (M) across different ECs (shown in the inset on the right × 10^12^/L) at specific ages.

#### Results based on erythrocyte count trajectories

The JM confirmed that males have higher EC than females. As estimated by the model, the average difference in EC between males and females is about 0.192 × 10^12^/L (variable “sex” in Supplementary Table S2, panel A). We also estimated a model including sex-time and sex-age interactions (Supplementary Table S3). According to this model, males have a faster decline in EC (∼ 0.011 × 10^12^/L per year) than females (p = 0.014), consistent with the empirical patterns shown in Figure 1 (Extended Data Figure 3).

Using the SPM (Methods and Supplementary Text), we found that at least one model component differed between females and males (p < 10^-16^). Based on this finding, we further determined which specific model components differed between the sexes. Consistent with the higher mortality rate in men, we found that the baseline mortality risk without the effect of EC is sex-dependent (p = 2.0 × 10^-4^). In addition, we confirmed sex dependence in model components reflecting the dynamics of ECs (p = 2.5 × 10^-3^) and their variability (p = 6.7 × 10^-4^). The component representing the estimated aging-related changes in ECs also showed sex dependence (p < 10^-16^). However, the ECT was not sex-dependent (p = 0.14).

We further investigated sex differences in the model component reflecting the estimated aging-related changes in ECs and found that both the intercepts and slopes of the respective trajectories were sex-dependent. Specifically, males exhibited a faster decline in the respective trajectory than females (by approximately 0.011 × 10¹²/L per year; p = 5.7 × 10⁻⁶), and the estimated initial value of this trajectory at age 70 years was higher in males than in females by approximately 0.33 × 10¹²/L (p < 10⁻¹⁶) (Extended Data Figure 4).

Figure 4 shows the relationships between EC decline with age, mortality risk, and survival in both sexes, based on the SPM. For instance, if a female has an EC = 4.4 × 10¹²/L at age 70 years (the average for her age) and her EC declines with age following the estimated average aging-related changes in ECs, then her EC will remain above the ECT (∼ 3.8 × 10¹²/L) at all ages shown in the figure. Consequently, her mortality risk will not differ from that of a female with any higher initial EC at age 70. However, for a female with an EC of 3.8 × 10¹²/L at age 70, the EC trajectory falls below the ECT, indicating an additional EC-specific mortality risk. For this female, the HR at age 90 years (relative to the mortality risk for a 70-year-old female with the average EC for her age) is 33.5, compared with an HR of 24.6 for a 90-year-old female whose EC at age 70 was average (inset table in the middle-left panel of Figure 4). For lower initial ECs, the differences in the respective HRs become even larger. Similar results are observed in males (middle-right panel).

**Figure 4.**
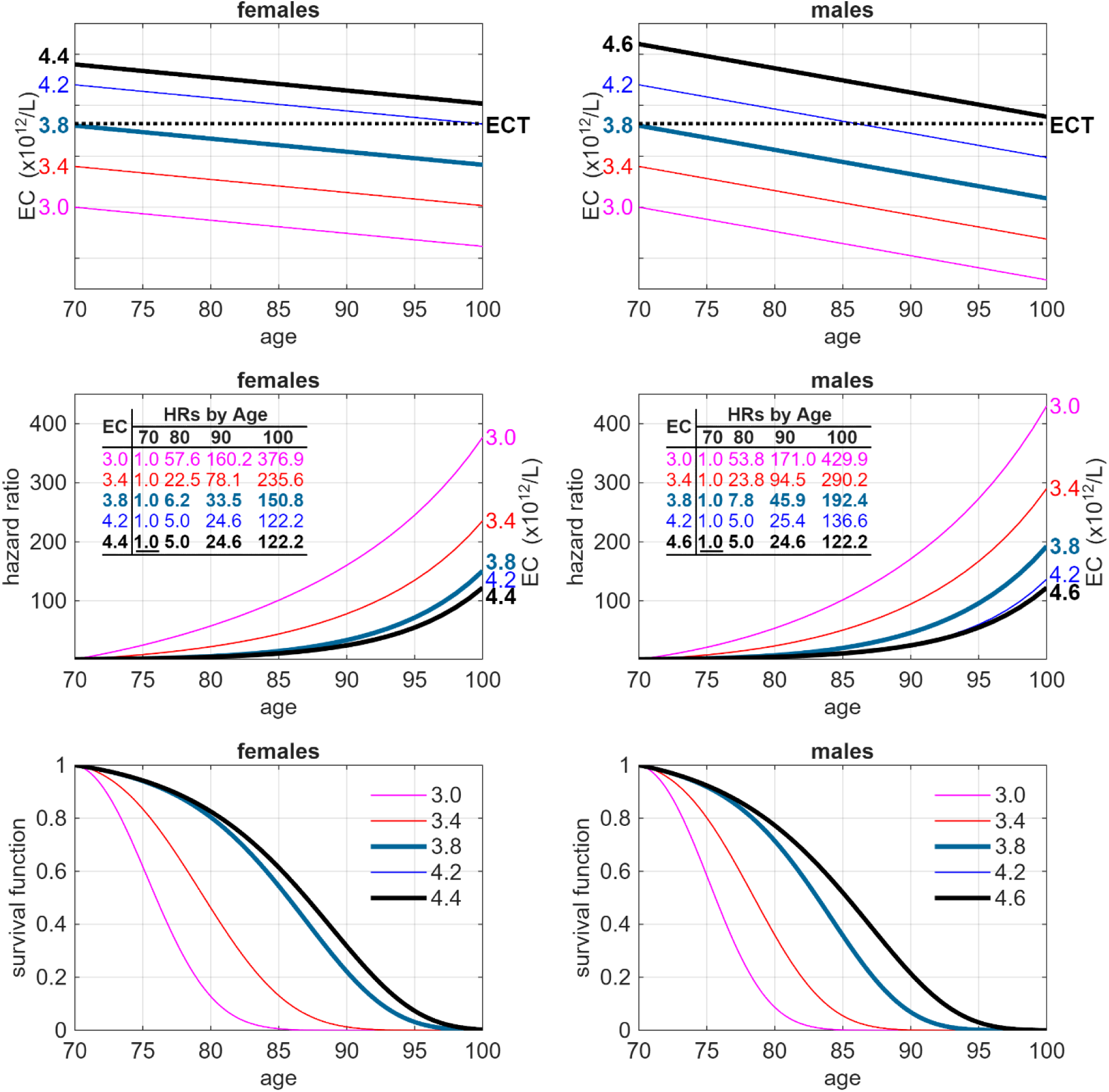
Associations between erythrocyte counts in females and males and mortality beyond age 70 years, based on the stochastic process model with an erythrocyte count threshold. The top panels show patterns of decline in erythrocyte count (EC) in females (top left) and males (top right) across different initial ECs at age 70 years. Dotted black lines depict the EC threshold (ECT). Middle panels display hazard ratios (HRs) estimated by the stochastic process model (Methods) for females (middle left) and males (middle right). The table presents numerical examples of HRs for specific ages and ECs, scaled to the reference age and EC (the reference HR is <u>underlined</u>). Bottom panels display survival curves (bottom left for females, bottom right for males) for different initial ECs (keys in the top-right corners) and the corresponding EC decline trajectories shown in top panels (e.g., the pink line denotes expected survivorship for individuals with an EC of 3 × 10^12^/L at 70 and respective decline with age). Note that the HR and survival curves for ECs of 4.2 and 4.4 × 10^12^/L in females overlap, and the survival curves for 4.2 and 4.6 × 10^12^/L in males are very close. Values above the reference HRs are not shown, as they would overlap with the reference HR curves. Two thick, bold lines correspond to the examples of initial ECs at age 70 years: 3.8 × 10^12^/L and average ECs at age 70 years (4.4 × 10^12^/L for females, 4.6 × 10^12^/L for males).

Based on the ECT, survival chances for individuals with ECs below the threshold are substantially lower than for those with ECs above it (Figure 4, bottom panels). For very low ECs, the differences in survival chances between females and males are small. However, at EC = 3.8 × 10¹²/L at age 70 years, a noticeable difference in survival emerges, because males exhibit a faster decline in EC. For example, a female with EC = 3.8 × 10¹²/L at age 70 years has about 54% (95% bootstrap CI: [31%, 65%]) chance of surviving to age 85 years, whereas a male with the same EC at that age has about a 36% (95% bootstrap CI: [14%, 55%]) chance of surviving to age 85 years.

Figure 5 displays the contributions of EC-independent and EC-dependent mortality to total mortality among female and male participants, as estimated by the SPM. Mortality above the ECT is similar across all ECs. However, for ECs below the ECT, the additional EC-dependent mortality increases overall mortality risk compared with levels above the ECT at each age. At older ages, this additional EC-related mortality increases more rapidly below the ECT than it does at younger ages.

**Figure 5.**
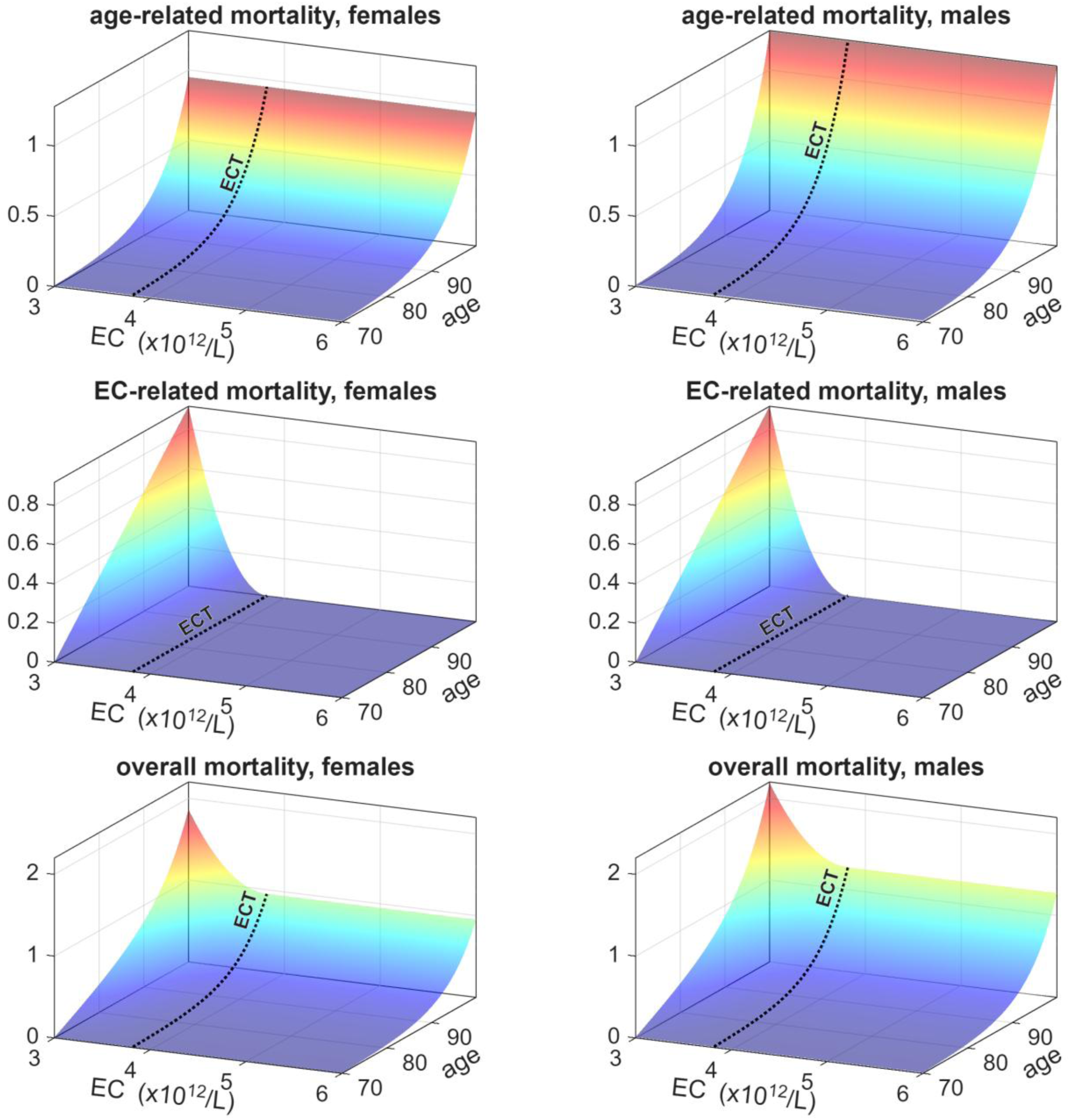
Mortality rates at different ages and erythrocyte counts in females and males, as estimated by the stochastic process model with an erythrocyte count threshold. The top panels show age-related (EC-independent) mortality rates. The middle panels show EC-related mortality rates. The bottom panels present the overall mortality rates. As specified by the model, overall mortality is the sum of age-related (EC-independent) mortality and additional EC-related mortality. The left panels show the rates for females, and the right panels show the rates for males. The dotted black line corresponds to the EC threshold (ECT). For ECs above the ECT, mortality risk does not depend on EC (i.e., additional EC-related mortality = 0 in the middle panels). For ECs below the ECT, an EC-related mortality amplifies overall mortality at a given age.

### Life span limits based on specific erythrocyte count trajectories, estimated by the SPM

Figure 6 presents the estimated ECT and the trajectories of aging-related changes in ECs, if these trajectories continue at the same rate at older ages beyond the maximum observed in the data. Although the ECT remains constant with age (p = 0.76), the estimated trajectory of aging-related changes in ECs declines progressively (p < 10⁻¹⁶). Consequently, if individuals live long enough, their ECs will eventually fall below the ECT (Figure 6). Under the fitted SPM, the estimated sex-specific trajectories of aging-related changes in ECs intersect the estimated threshold at approximately 126 years in females and 114 years in males. These values should be interpreted as model-based extrapolations derived from point estimates of model parameters, rather than as direct estimates of lifespan limits. Importantly, EC-associated mortality would not occur in many participants because their ECs remain above the ECT even at very advanced ages. For example, ECs below the ECT were observed in 22.7% (95% bootstrap CI: [5.7%, 36.6%]) of participants aged 90 years or older and 36.5% (95% bootstrap CI: [11.1%, 52.4%]) of those aged 100 years or older.

**Figure 6.**
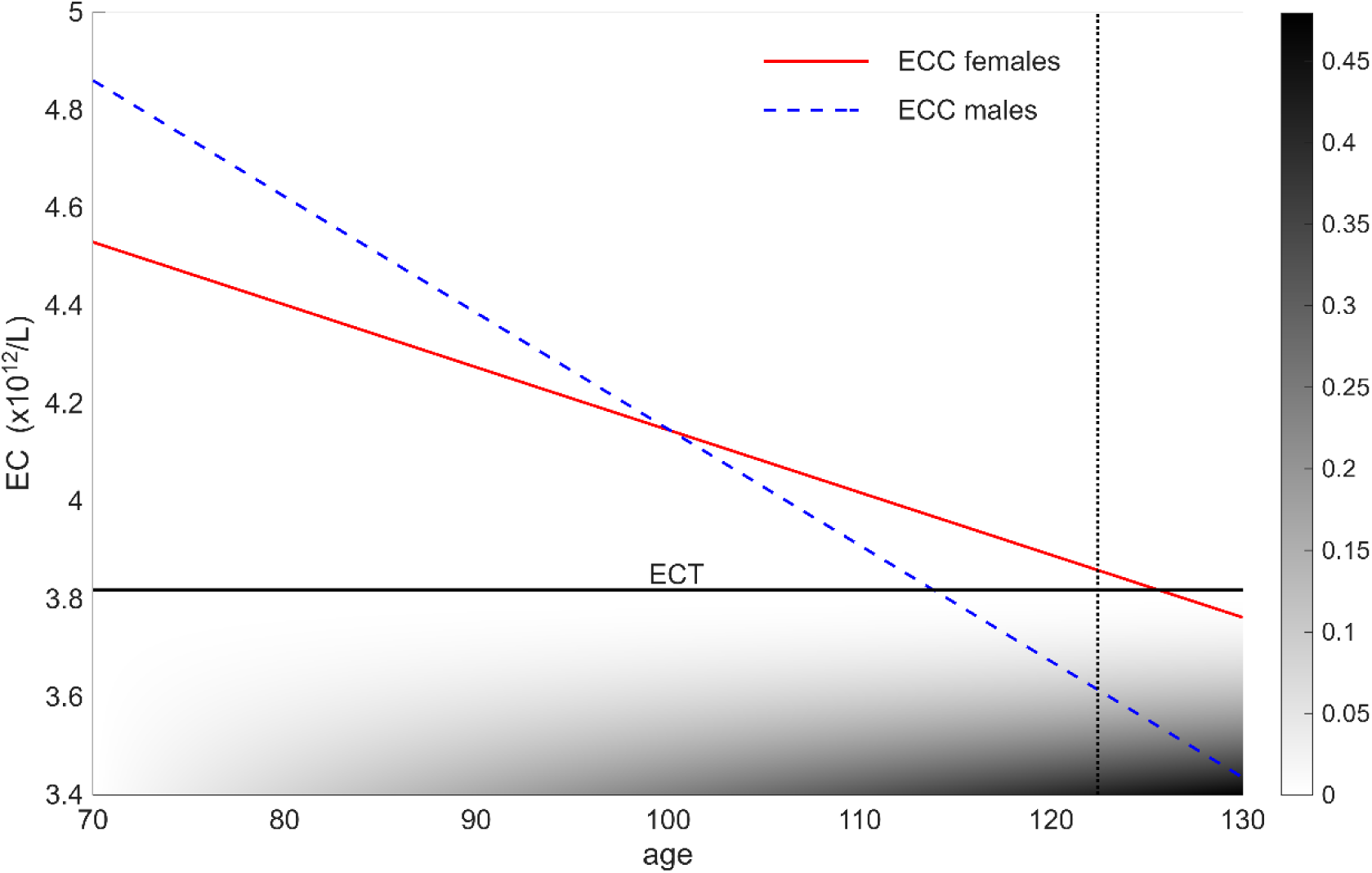
Trajectories of aging-related changes in erythrocyte count and the erythrocyte count threshold, estimated by the stochastic process model. The solid red and dashed blue lines correspond to the estimated aging-related erythrocyte count (EC) changes (ECC) for females and males, respectively. The solid black line depicts the EC threshold (ECT) that triggers additional EC-associated mortality, as estimated by the model. The grey shading below the ECT denotes EC-associated mortality (see the scale on the right). The vertical black dotted line denotes the longest documented human lifespan (122 years and 164 days ^44^). Note that although the ECC trajectories differ between females and males (p = 5.7 × 10^-6^), the ECT is the same for both sexes (p = 0.14). Methods and Supplementary Text provide additional details on the model.

**Figure 7.**
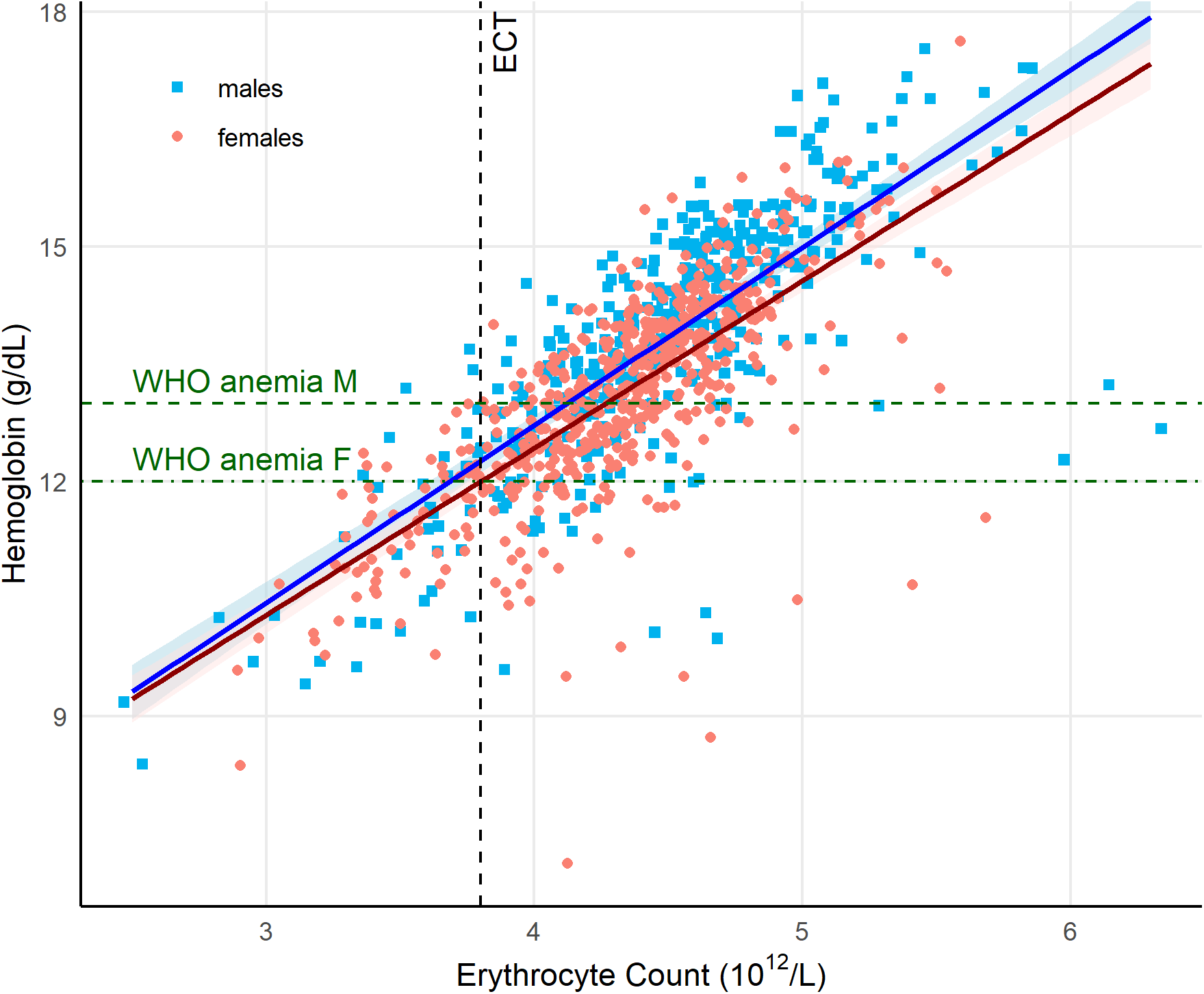
Relationships between baseline erythrocyte counts and hemoglobin. Regression lines (females, dark red; males, blue) display hemoglobin (Hgb) for different erythrocyte counts (ECs), as obtained from the fixed-effects component of a linear mixed-effects model adjusted for age (Methods). Predictions were generated across a grid of ECs spanning the observed range in the analytic sample (95% confidence intervals for the predicted values are displayed as shaded bands around regression lines). A vertical dashed black line indicates the EC threshold (ECT = 3.8 × 10¹²/L) estimated from the stochastic process model (Methods). Horizontal dashed green lines denote WHO anemia thresholds: 13 g/dL for males (dashed line labeled “WHO anemia M”) and 12 g/dL for females (dashed line labeled “WHO anemia F”). Observed values for females and males are shown as pink circles and blue squares, respectively. Horizontal jittering was applied to ECs for visualization.

### Sensitivity analyses

Parameter estimates generated by the CPHM were adjusted for additional covariates representing factors that may affect EC or mortality, e.g., inflammation and iron deficiency anemia (Methods). This expanded model confirmed the association between lower ECs and increased mortality risk: HR per 1 × 10^12^/L decrease in EC increased from 1.32 (without additional covariates) to 1.37 (95% bootstrap CI: [1.13, 1.68]) (Supplementary Table S4). We further tested this association by applying the CPHM to samples with older age cutoffs (75, 80, 85, and 90 years). The association between lower ECs and increased mortality risk remained significant across these older age groups, despite smaller sample sizes (Supplementary Table S5).

We included the same additional covariates in the JM. As in the analyses of baseline EC measurements, the association between lower EC trajectories and increased mortality risk remained significant in this expanded model (Supplementary Table S6). The estimated regression coefficient for EC trajectories in this model was -0.333, corresponding to an HR of 1.4 (95% bootstrap CI: [1.09, 1.79]) for a unit (1 × 10^12^/L) decrease in EC trajectories below the expected ECs (variable “EC_ind_”, Supplementary Table S6, panel B). The expanded model also confirmed that males have higher EC than females (variable “sex”, Supplementary Table S6, panel A) and that males exhibit a faster decline in EC (variable “sex*time”, Supplementary Table S6, panel A).

### The relation between the erythrocyte count threshold and the WHO anemia definition

The WHO defines anemia as Hgb below thresholds of 13 g/dL in males and 12 g/dL in females. We thus examined the relation between the ECT and these WHO thresholds using mixed models applied to baseline EC and Hgb from our analytic sample (Methods). Based on the ECT, the proportions of participants at increased mortality risk (i.e., below the ECT) changed substantially with shifting the Hgb threshold from Hgb <13 g/dL to Hgb <12 g/dL, both in the entire sample (25% vs. 47%, p=1.2 × 10^-6^) as well as in sex-specific samples (females: 23% vs. 45%, p=3.4 × 10^-5^; males: 29% vs. 51%, p=0.01). In an extended multivariable-adjusted mixed model, which included covariates reflecting iron reserves, inflammation, and kidney function (Extended Data Figure 5), there was no significant change in the slopes of the regressions between Hgb and ECs.

## DISCUSSION

LLFS participants exhibited an age-related decline in ECs, evident in both baseline and sequential measurements. Although males had higher ECs than females at age 70 years, their age-related decline in ECs was steeper. Analysis with CPHM and the JM showed that lower ECs were associated with an increased risk of mortality based on both baseline and sequential EC measurements. Analysis with the SPM further revealed a sharp increase in mortality risk and, correspondingly, a marked decline in survival when ECs fell below an estimated ECT of 3.8 × 10¹²/L.

Based on the WHO definition, the prevalence of anemia increases after age 60 years ^6–9^, and exceeds 50% in centenarians ^13,14^. To what extent does the age-related decline in ECs contribute to anemia and its associated outcomes in older adults? Approximately one-third of anemias in adults aged 60 years or older are attributable to deficiencies in essential nutrients, particularly iron, and another one-third to chronic diseases, including renal disease, cancer, heart disease, and inflammatory disorders ^6^. These conditions are collectively referred to as explained anemia of aging (EAA). Still, in one-third of older adults with anemia, the causes remain unidentified despite comprehensive clinical evaluation. This condition is known as unexplained anemia of aging (UAA) ^6,9,15^.

However, the ECT findings, derived from computational and modeling analyses of mortality risk associated with the progressive decline in EC among LLFS participants ≥70 years old, raise questions about extending the WHO anemia definitions, based on populations ≤65 years old ^10,11^, to both EAA and UAA. These forms of anemia that emerge in older adults with progressively declining EC are unlikely to share major causes or consequences with anemia diagnosed in children and younger adults. The discordance between the sex-specific WHO anemia thresholds and the sex-independent ECT further supports this distinction.

Based on the ECT, many LLFS participants who met WHO anemia criteria had similar mortality risk, with or without adjustment for covariates commonly associated with EAA, including iron stores, chronic inflammation, and renal function. Thus, the progressive decline in ECs in older adults may contribute to both EAA and UAA, as well as to their associated mortality. That is, if an individual’s ECs are constrained by intrinsic factors of aging, lower ECs due to nutritional deficiencies, for instance, may lead to even worse anemia than they would otherwise. Such a conclusion raises a fundamental question about the core etiology of not only UAA—a topic of interest to the National Institute on Aging ^16^ —but also of EAA.

A progressive decline in EC with age may result from reduced survival of circulating erythrocytes, diminished erythropoiesis, or both. While empirical evidence on age-related changes in erythrocyte survival is scarce, a mid-20th-century chromium-labeling study on a small number of individuals suggests that the erythrocyte lifespan is largely preserved during healthy aging ^17^. Thus, reduced erythropoiesis is likely the cause of the progressive decline in EC in older adults and the development of anemia.

Anemia, including EAA and UAA, is associated not only with higher mortality but also with declines in physical and cognitive function ^15,18,19^. In recent years, extensive research on the biology of aging has identified key processes that drive aging, collectively termed the hallmarks of aging ^20^. Among these are processes considered “primary” drivers, including genomic instability, epigenetic alterations, proteostasis loss, and telomere shortening.

Telomere biology likely contributes to EAA and UAA in older adults. First, the discovery by Hayflick and Moorhead ^2^ indicated that human somatic cells possess a built-in replication counter. A decade later, this counter was identified as telomere shortening with each cycle of cell replication, which eventually produces critically short telomeres incapable of sustaining further replication ^21^. Second, because erythropoiesis is the most intensive replicative process in the human body ^4^, telomere shortening in hematopoietic stem cells, as reflected in leukocyte telomere shortening with age ^22,23^, proceeds more rapidly than in other somatic cells. As a result, leukocytes have the shortest telomeres among adult somatic cell types ^24^. Third, humans have short telomeres and long lifespans ^25^, suggesting that telomere length may constrain EC production with age. This likely contributes to an age-related decline in erythropoiesis, consistent with the observed correlation between EC and leukocyte telomere length in adults ^26,27^. Furthermore, leukocyte telomere length in about 50% of centenarians ^28^ is as short as that seen in patients with telomere biology disorders ^29^, which result from deleterious mutations in telomere maintenance genes. These patients often develop anemia and die young. Collectively, these observations suggest that by diminishing erythropoietic reserve and increasing vulnerability to anemia-associated morbidity and mortality, short telomeres may affect survival to exceptionally old age in many humans.

We note the strengths of our analytical approach. First, we applied three complementary analytic approaches, CPHM, JM, and SPM, to evaluate the association between EC and mortality from several perspectives, including baseline EC measurements, longitudinal EC trajectories with age, and the ECT. Second, we analyzed a unique cohort from the LLFS enriched for families with exceptional longevity. This cohort is particularly well-suited for examining the relationship of EC and anemias of aging with mortality in individuals approaching their NLS limit.

The limitations of the work should also be considered. The LLFS comprises predominantly White participants from families prone to exceptional longevity, and it is known to be a healthy aging cohort ^30^. Nonetheless, these findings provide a strong foundation for testing their generalizability across broader, more diverse populations. Moreover, the observed association between EC and mortality does not imply causality, and SPM-based survival extrapolations in LLFS participants are not direct estimates of human lifespan limits. The ECT reported in this work was derived from an SPM that included only a limited set of covariates (age and sex).

Exploring additional covariates in the SPM would require larger sample sizes. Finally, the proposed link between hematopoietic telomere biology, EC dynamics, and the NLS limit should be regarded as speculative.

In conclusion, demographers have rarely considered natural constraints on human lifespan. Although such constraints are likely multifactorial, our findings suggest that the age-related decline in EC, manifested as anemias of aging, amplifies the exponential increase in mortality and may represent one such constraint. EAA and UAA should therefore be defined by health and longevity outcomes rather than Hgb cutoffs derived from younger populations. Extending lifespan beyond its current limit may depend, in part, on preserving erythropoietic capacity with age.

## ACKNOWLEDGEMENTS

This research is supported by NIH grant 2U19AG063893-06 “The Long Life Family Study”. AA’s recent telomere research has been supported by NIH grants U01AG066529 and R01ES035760.

## AUTHOR CONTRIBUTIONS

KGA and AA designed the study and were the primary writers of the manuscript. KGA and OB performed all modeling and statistical analyses. JMM, HJC, WEJ, and DTAE provided guidance for further analyses and contributed to manuscript writing. BLP provided advice regarding the telomere-erythrocyte count connection. SL advised on joint model analyses and contributed to manuscript writing. AIY provided advice on stochastic process model analyses and contributed to manuscript writing. MCM, SLA, KC, JMZ, BT, and MAP contributed to manuscript writing.

## ONLINE METHODS

### Participant characteristics and data acquisition

The Long Life Family Study (LLFS) is a longitudinal, multigenerational family study of healthy aging and exceptional longevity ^12^. Between 2006 and 2009 (Visit 1), LLFS recruited 539 families selected for familial longevity, comprising 4,953 family members and spouses.

Participants were enrolled through three US field centers (Boston University, Boston, MA; Columbia University, New York, NY; and the University of Pittsburgh, Pittsburgh, PA) and the University of Southern Denmark, Odense, Denmark. Participants were selected from the upper 1% tail of the Family Longevity Selection Score ^31^ distribution, which quantifies exceptional familial longevity.

The study includes detailed in-person evaluations at three visits. Visit 1 included long-lived individuals (probands), their siblings, spouses, and offspring, as well as spouses of the offspring. Visit 2 (2014–2017) included surviving participants from Visit 1 and newly enrolled participants (i.e., additional family members from the two generations of the existing families). Visit 3 (2020-2024) includes surviving participants from Visits 1 and 2, as well as newly enrolled participants, including grandchildren (Third Generation). Participants provided information on socio-demographic indicators, past and current medical conditions, medication use, and physical and cognitive functioning ^12^. Blood samples were collected on each visit. Unprocessed ethylenediaminetetraacetic acid (EDTA) tubes were shipped to the Advanced Diagnostics and Research Laboratory (ARDL) at the University of Minnesota for complete blood count measurements. Additional technical details on biospecimen collection procedures in the LLFS can be found in ref. ^32^ and on the study website (https://longlifefamilystudy.com/manual-of-procedures/).

We used the August 19, 2024 release of the LLFS data. Baseline ages were validated using dates of birth from official documents (e.g., driver’s licenses, birth certificates) in the U.S. ^33^ and through the Danish civil registration system. Vital status was ascertained for all participants at each annual follow-up contact ^34^. We calculated ages at censoring for living participants using birth dates and the most recent follow-up dates. We used erythrocyte count (EC) measurements from individuals aged 70 years and older. The 70-year age cutoff was selected because relatively few deaths occur at younger ages in the LLFS, limiting statistical power in those age ranges (see also the description of sensitivity analyses using different age cutoffs in the next section). Participants with cancer or kidney disease were excluded (1537 or 49.4% of all eligible individuals aged 70 years and older), since both conditions and cancer treatments can influence EC. These exclusions applied to both cross-sectional and longitudinal empirical analyses, as well as to statistical modeling. Extended Data Figure 1 shows a flowchart describing how the analytic sample was constructed from the original LLFS dataset. Table 1 presents the characteristics of the analytic sample used in the main analyses reported in the paper. Because all statistical analyses were performed using complete-case samples, the sensitivity analyses described below used smaller subsamples with non-missing additional covariates, as reported in Supplementary Table S5 and in the text.

### Statistical analysis: First erythrocyte count measurement at age 70 years or older, and mortality risk

We investigated the association between EC and mortality among LLFS participants using the Cox proportional hazards model (CPHM) implemented in the *survival* R package (version 3.8-6). The analysis used the first EC measurement at age 70 years or older as the primary predictor of mortality risk, time since that observation as the time scale, and mortality (follow-up time until death or censoring, with an indicator: 1 – died, 0 – censored) as the time-to-event outcome. The main model was adjusted for age at enrollment and sex (1 – male, 0 – female). Age was centered at 70 years, and EC was centered at 4.5 × 10^12^/L. The proportional hazards assumption was tested using the *cox.zph* function from the *survival* package and was confirmed (p > 0.05) for the main variable (EC) and both covariates. Cumulative baseline hazards were extracted from the fitted Cox model using the *basehaz* function with option *centered = FALSE*. The cumulative baseline hazards were then fitted using a Gompertz function to obtain smooth curves, but the Gompertz fit was not used in the regression inference. These fitted curves, together with the estimated model parameters, were used only to generate female and male survival curves for different ECs at age 70, as presented in the main text.

In additional sensitivity analyses, we performed a robustness check using the model further adjusted for country (1 – Denmark, 0 – USA) and for additional covariates, including smoking, body mass index (BMI), cystatin C (CysC), high sensitivity C-reactive protein (hsCRP), interleukin-6 (IL-6), and transferrin receptor (sTfR). These covariates reflect health status factors that can affect EC or mortality, e.g., inflammation and iron-deficiency anemia. Continuous variables (BMI, CysC, hsCRP, IL-6, and sTfR) were inverse-normally transformed for the analyses. We also applied this model to other cut-off ages (75 years and older [75+], 80+, 85+, and 90+ years). When testing the proportional hazards assumption in these expanded models, we found violations in several instances for age, IL-6, and BMI. To address this, we used binary variables IL6bin (1 – IL-6 above median, 0 – IL-6 below median) and BMIbin (1 – BMI above median, 0 – BMI below median) and included age as a stratification variable (with age groups 70-74, 75-79, 80-84, 85-89, and 90+) in the expanded models.

Sandwich variance estimators clustered by family/pedigree were used to account for relatedness in the Cox model. We also used the familial bootstrap approach ^35^ to produce 95% confidence intervals, estimated using the percentile method, for the hazard ratios and other derived quantities (e.g., survival probabilities) reported in the text.

### Statistical analysis: Longitudinal erythrocyte count measurements and mortality risk

We modeled the joint evolution of longitudinal EC trajectories and time-to-death among LLFS participants aged 70 years or older using the shared-random-effects joint model (JM) framework as implemented in the *joineRML* R package (version 0.4.7) ^36,37^. The model was applied to the same sample used in the CPHM analysis, i.e., individuals with at least one EC measurement at ages 70 years or older. In the main model, we used the same covariates (age and sex) in both the survival and longitudinal submodels as in the CPHM. In sensitivity analyses, we estimated an expanded model that included additional covariates (country, smoking, BMI, CysC, hsCRP, IL-6, and sTfR), paralleling the CPHM specification. For the analyses reported in “The sex effect,” we also estimated a model that included sex-by-age and sex-by-time interactions in the longitudinal submodel. Time since baseline was used as the time scale.

The longitudinal submodel used linear random effects (random intercept and slope), and the survival submodel was a Cox-type proportional hazards model with an unspecified baseline hazard, linked to the longitudinal process through a single scalar association parameter, as implemented in *joineRML*. Specifically, for subject *i* (*i* = 1, …, *n*), with EC measured at times *t_ij_*, the longitudinal process was modeled as follows:

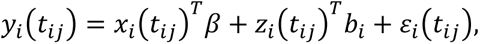

where *ε_i_*(*t_ij_*)∼*N*(0, *σ*^2^) are independent measurement errors; the vector *b_i_* contains the subject-specific random effects (random intercept and slope in our applications, corresponding to *z_i_*(*t*)*^T^* = (1, *t*)*^T^*, with superscript *T* denoting transposition), assumed to follow a multivariate normal distribution *b_i_*∼*N*(0, *D*). The matrix *D* is the variance–covariance matrix of the random effects: it quantifies between-subject variability in baseline EC (random intercept), variability in the rate of change in EC (random slope), and their covariance. Its diagonal elements represent the variances of the random intercept and random slope, and its off-diagonal element represents their covariance, capturing the association between a subject’s baseline EC and the rate of change in EC. The fixed-effects design vector *x_i_*(*t_ij_*) contains the covariates included in the model, and the corresponding regression parameters are represented by the vector *β*.

For the survival submodel, we specified the hazard at time *t*, ℎ*_i_*(*t*|*b_i_*), as a proportional hazards model with an unspecified baseline hazard ℎ_0_(*t*):

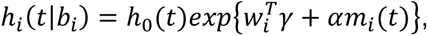

where *w_i_* includes baseline covariates, *γ* is the corresponding vector of regression coefficients, and *m_i_*(*t*) = *z_i_*(*t*)*^T^b_i_* (denoted EC_ind_, Supplementary Tables S2, S3, and S6) is the subject-specific random-effects linear predictor at time *t* (i.e., random intercept plus *t* times random slope). The parameter *α* is a scalar association coefficient that links the deviation of an individual’s EC trajectory from the population mean to the hazard of death.

In the longitudinal submodel, the expected (“population mean”) EC for subject *i* at time *t* is *E*[*y_i_*(*t*)] = *x_i_*(*t*)*^T^β*. The term *m_i_*(*t*) = *z_i_*(*t*)*^T^b_i_* represents the subject-specific deviation from this expected EC due to the individual’s random intercept and slope. Thus, *m_i_*(*t*) > 0 indicates that the individual’s EC is higher than expected based on the covariates and population averages at time *t*; whereas *m_i_*(*t*) < 0 indicates it is lower.

The coefficient *α* quantifies how subject-specific deviations in EC (*m_i_*(*t*)) are associated with mortality. Because *joineRML* implements a proportional hazards model for the survival submodel, the interpretation of *α* and the corresponding hazard ratio (HR) is analogous to that in a Cox regression model. A one-unit increase in *m_i_*(*t*) multiplies the hazard at time *t* by *HR_α_* = *exp*(*α*). A negative *α* indicates that being above the population-mean EC trajectory (i.e., *m_i_*(*t*) > 0) is protective and reduces the hazard. For example, suppose *α* = -0.35, then the instantaneous hazard ratio for a one-unit increase in *m_i_*(*t*) is *HR_α_* = *exp*(−0.35) ≈ 0.7. Thus, consider two individuals with the same covariates. At a given time *t*, individual 1 has an EC equal to the population mean (*m_i_*(*t*) = 0), while individual 2 has an EC one unit higher on the longitudinal submodel’s scale (i.e., 10^12^/L in our case, so *m_i_*(*t*) = 1). In this scenario, individual 2 has a 30% lower hazard at time *t* compared with individual 1.

As in the CPHM analyses, we used the familial bootstrap approach ^35^ to produce 95% confidence intervals for the regression coefficients and HRs, which were reported alongside the original *joineRML* outputs in Supplementary Tables S2, S3, and S6, as well as in the main text.

### Statistical analysis: Testing the erythrocyte count threshold hypothesis using the stochastic process model

We applied the stochastic process model (SPM) ^38,39^ to analyze the association between longitudinal ECs and mortality among LLFS participants aged 70 years or older. The SPM is particularly relevant in aging research because it combines the statistical rigor of general methodology with biologically grounded assumptions ^38–41^. Modeling EC using stochastic differential equations allows representation of both systematic drift (e.g., homeostatic control or “adaptive capacity”) and random fluctuations (diffusion) that contribute to observed variability. When integrated with time-to-event outcomes such as mortality, these models can quantify how deviations from “optimal” trajectories translate into risk. Technical details of the modified SPM implemented in our analysis are provided in the Supplementary Text. A non-technical description of the general SPM can be found in our earlier paper ^41^. Additional information on the interpretation and illustration of model components, parameters, and related null hypotheses is presented in the Supplementary Materials of our recent publication ^40^.

In brief, the SPM is a versatile model for the joint analysis of longitudinal and time-to-event outcomes, developed for aging research. It shares key advantages with generic joint models (e.g., the ability to accommodate missing-not-at-random mechanisms common in aging studies), while offering additional flexibility relevant to this field. In particular, the SPM allows the general association between longitudinal and time-to-event outcomes (which can be estimated using a JM) to be decomposed into components representing specific aging-related characteristics embedded in the model, enabling hypothesis testing about their age patterns and relationships to variables of interest ^40^.

In our application to EC trajectories and mortality, we hypothesize the existence of an EC threshold (ECT), which may vary with age, such that mortality risk is unaffected by ECs above the ECT, but when EC falls below the threshold, an additional mortality component is activated; this results in mortality exceeding the baseline level at the corresponding age. This additional risk is modeled as a quadratic function of EC (for a given age).

The SPM also models an “equilibrium” trajectory, that is, the long-term mean to which the EC trajectory converges, as the equilibrium EC. In general, this equilibrium trajectory may differ from the ECT, and both may depend on age or other variables of interest (tests for such dependencies are incorporated in the model). In addition, the model allows testing whether the quadratic shape of the additional mortality component narrows with age (i.e., the same deviation of EC from the ECT induces a larger mortality increase at older ages) and whether other covariates (e.g., sex) influence the width of this EC-related mortality component.

Following our earlier applications of the SPM to LLFS data ^40^ and the typical age pattern of human mortality at older ages, we used a Gompertz baseline mortality function (representing mortality unrelated to EC), adjusted for sex. Other model components were assumed to be linear functions of age and sex, with two exceptions: the EC-related mortality multiplier, which was assumed to be the same for females and males, and the variability parameter, which was assumed to be age-independent (Supplementary Text). This parsimonious specification allows testing hypotheses about the dependence of model components on age and sex. We tested the key hypothesis regarding the existence of an ECT, that is, that a non-zero additional mortality term appears when ECs fall below the ECT.

We also tested other null hypotheses as described in the Supplementary Text. These hypotheses were evaluated using the likelihood ratio tests, as in our earlier applications ^40^. As with the JM, we applied the SPM to individuals with at least one EC measurement at ages 70 years or older (noting that individuals with only one observation still contribute to the likelihood function, thereby increasing statistical power ^42,43^). We used the familial bootstrap approach ^35^ to generate 95% confidence intervals for the model parameters. Point estimates and 95% familial bootstrap confidence intervals, calculated using the percentile method, are reported in the main text. The SPM analysis was performed using in-house MATLAB code run in version R2024a.

**Statistical analysis: Relationships between baseline erythrocyte counts and hemoglobin concentration.** The WHO defines anemia as hemoglobin (Hgb) concentrations below thresholds of 13 g/dL in males and 12 g/dL in females ^10^. We examined the relationship between the ECT, as estimated using the SPM described above, and these WHO thresholds using baseline EC and Hgb from our analytic sample. To account for familial clustering in LLFS, linear mixed-effects models were fitted with a random intercept for family, with baseline Hgb as the outcome and baseline EC as the primary predictor. The fixed-effects component also included baseline age in a basic model reported in the main text, and additional adjustments for country, smoking, body mass index, cystatin C, high-sensitivity C-reactive protein, interleukin-6, and transferrin receptor in an extended model reported in the Extended Data. The models were fitted separately for males and females. All continuous covariates (except EC and Hgb) were centered at their sample means to facilitate interpretation.

## EXTENDED DATA FIGURES

**Extended Data Figure 1.**
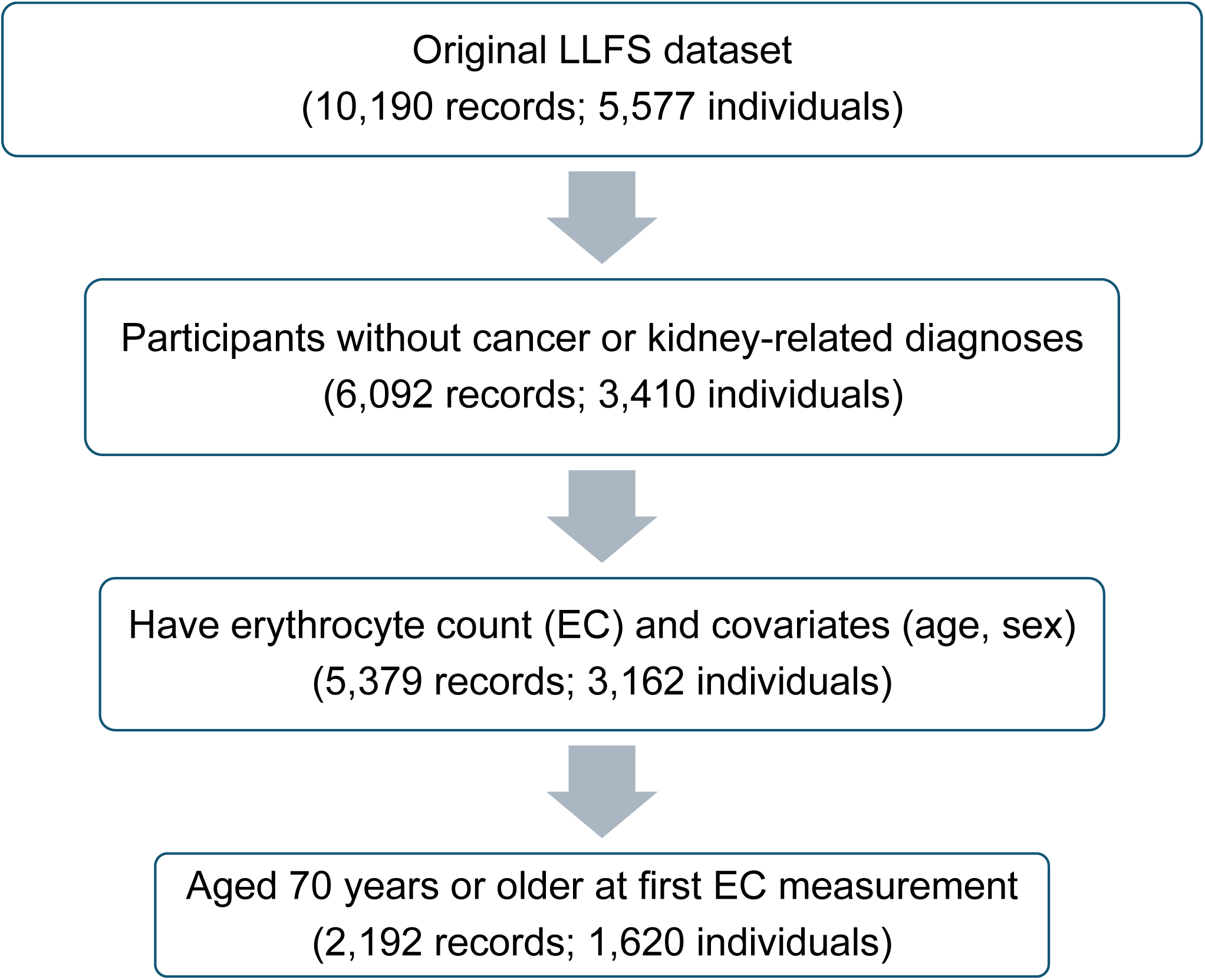
Flowchart, illustrating the selection of LLFS participants for analysis.

**Extended Data Figure 2.**
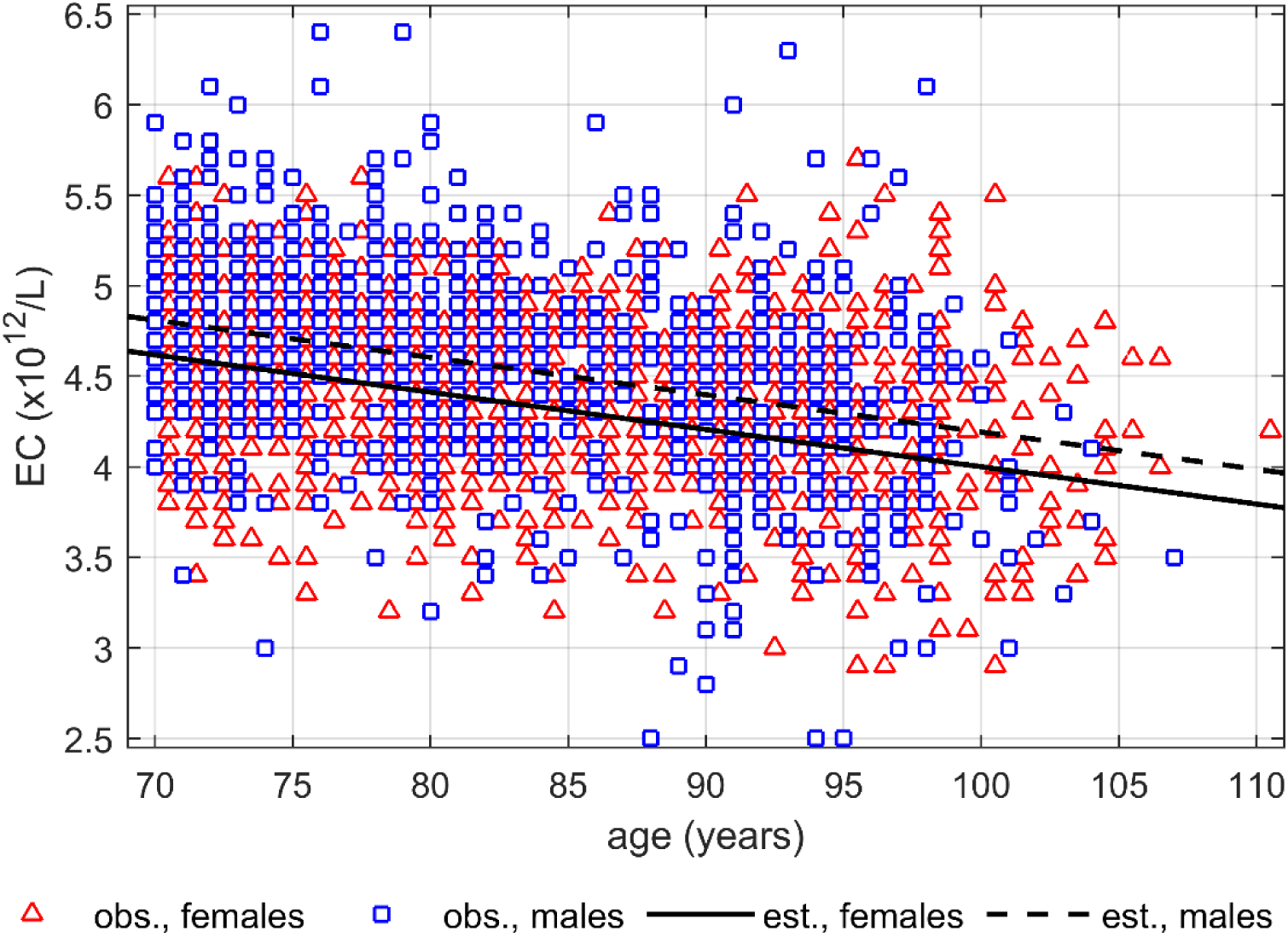
Average age trajectories of erythrocyte counts (ECs) in females and males estimated from the joint model, adjusted for basic demographic characteristics, and applied to data on mortality and ECs in LLFS participants aged 70 years or older. Estimated trajectories are shown as black lines (solid for females, dashed for males). Open red triangles and blue squares display the ECs for females and males, respectively. Females’ values were jittered horizontally for better visibility.

**Extended Data Figure 3.**
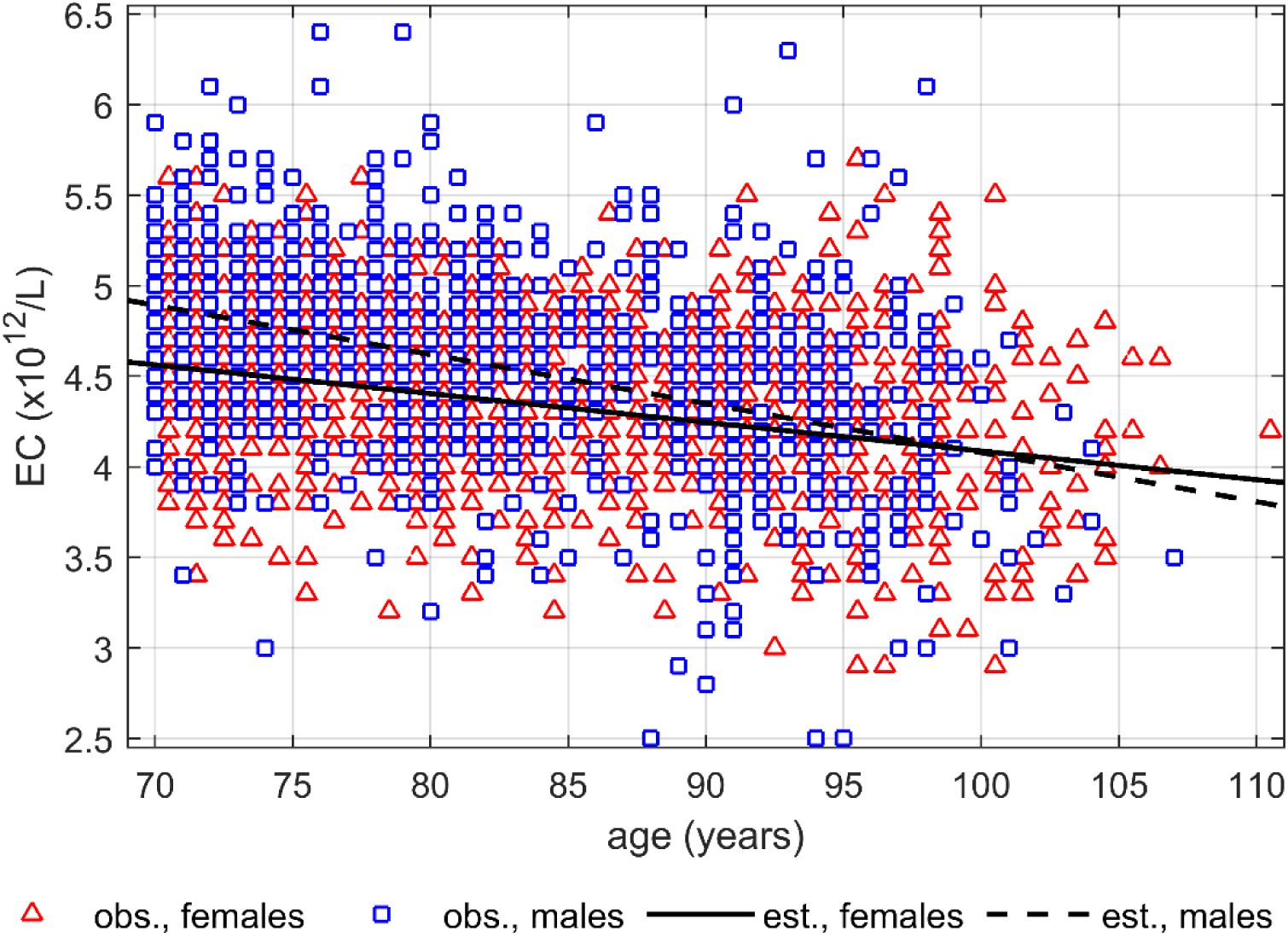
Average age trajectories of erythrocyte counts (ECs) in females and males estimated from the joint model, adjusted for basic demographic characteristics and age-time and sex-time interaction terms, and applied to data on mortality and ECs in LLFS participants aged 70 years or older. Estimated trajectories are shown as black lines (solid for females, dashed for males). Open red triangles and blue squares display the ECs for females and males, respectively. Females’ values were jittered horizontally for better visibility.

**Extended Data Figure 4.**
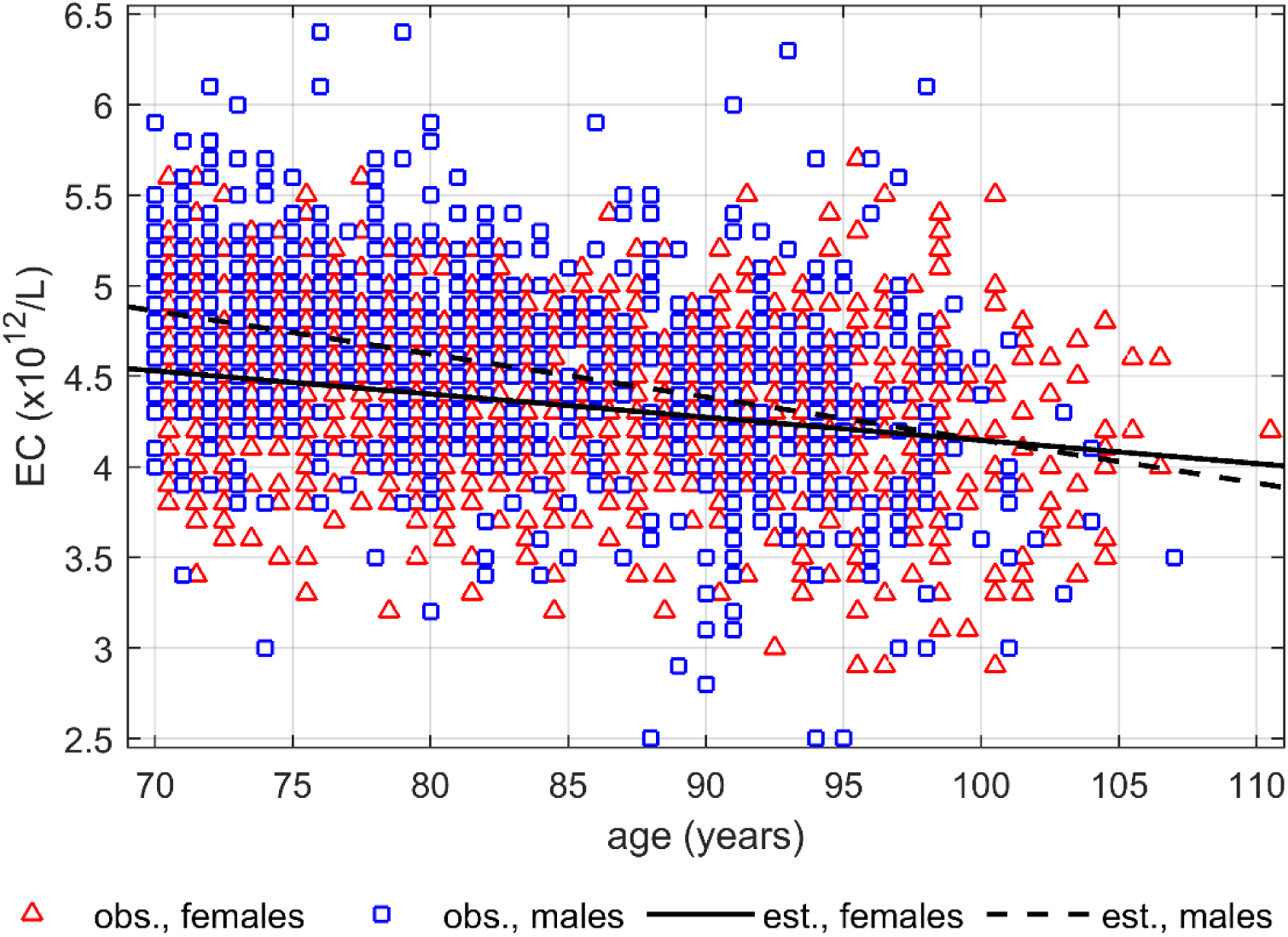
Estimated aging-related changes in erythrocyte counts (ECs) based on the stochastic process model applied to data on mortality and ECs in LLFS participants aged 70 years or older. Estimated trajectories are shown as black lines (solid for females, dashed for males). Open red triangles and blue squares display the ECs for females and males, respectively. Females’ values were jittered horizontally for better visibility.

**Extended Data Figure 5.**
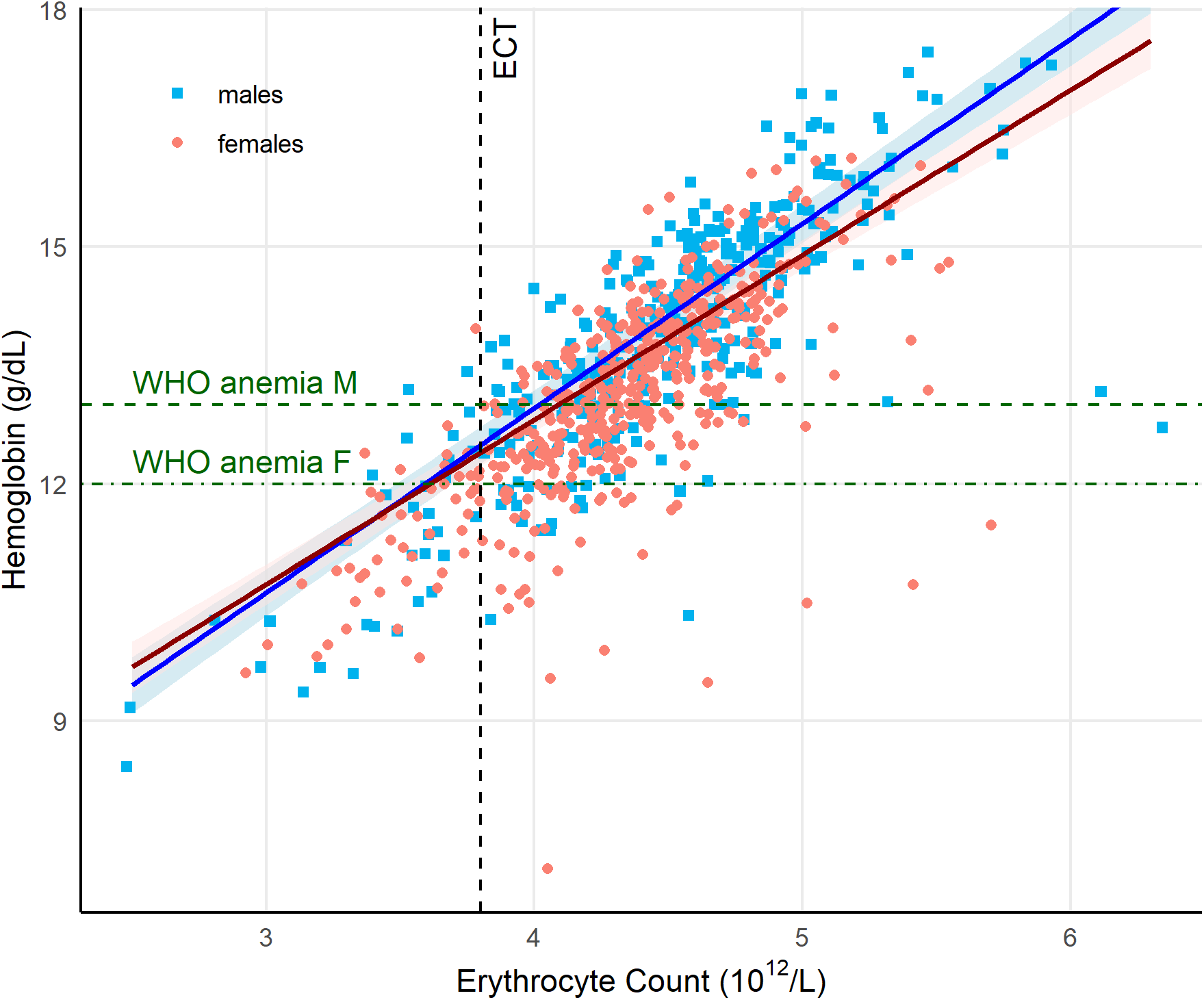
Relationships between baseline erythrocyte counts and hemoglobin in the extended model. Regression lines (females, dark red; males, blue) display hemoglobin (Hgb) across different erythrocyte counts (ECs), as obtained from the fixed-effects component of a linear mixed-effects model adjusted for age, country, smoking, body mass index, cystatin C, high-sensitivity C-reactive protein, interleukin-6, and transferrin receptor (Methods). Predictions were generated across a grid of ECs spanning the observed range in the analytic sample (95% confidence intervals for the predicted values are displayed as shaded bands around regression lines). A vertical dashed black line indicates the EC threshold (ECT = 3.8 × 10¹²/L) estimated from the stochastic process model (Methods). Horizontal dashed green lines denote WHO anemia thresholds: 13 g/dL for males (dashed line labeled “WHO anemia M”) and 12 g/dL for females (dash-dotted line labeled “WHO anemia F”). Observed values for females and males are shown as pink circles and blue squares, respectively. Horizontal jittering was applied to ECs for visualization.

