## Supplementary Materials for "Erythrocyte Count, Anemia, and the Human Natural Lifespan Limit: Evidence from the Long Life Family Study"

### SUPPLEMENTARY INFORMATION

#### Inventory of Supplementary Information

**Supplementary Text.** Technical description of the stochastic process model

**Supplementary Table S1.** Detailed characteristics of the Long Life Family Study samples with one or more erythrocyte count (EC) measurements (EC1+) and two or more EC measurements (EC2+) used in the analyses

**Supplementary Table S2.** Results of the joint model adjusted for basic demographic characteristics, applied to data on mortality and erythrocyte counts in LLFS participants aged 70 years or older

**Supplementary Table S3.** Results of the joint model adjusted for basic demographic characteristics and age-time and sex-time interaction terms, applied to data on mortality and erythrocyte counts in LLFS participants aged 70 years and older

**Supplementary Table S4.** Results of the Cox proportional hazards model adjusted for additional covariates, applied to data on mortality and the first observation of erythrocyte counts in LLFS participants aged 70 years and older

**Supplementary Table S5.** Results of the Cox proportional hazards model adjusted for basic demographic characteristics and additional covariates, applied to data on mortality and the first observation of erythrocyte counts in LLFS participants from different age groups

**Supplementary Table S6.** Results of the joint model adjusted for basic demographic characteristics and additional covariates, applied to data on mortality and erythrocyte counts in LLFS participants aged 70 years and older

### Supplementary Text. Technical description of the stochastic process model

We extended the basic stochastic process model (SPM) [1,2] by incorporating an erythrocyte count (EC) threshold (ECT), such that there is no association between EC and mortality when ECs are above the ECT. Specifically, the modified SPM consists of two equations: one describing the dynamics of the repeatedly measured variable (EC) for individual *i* (denoted $Y_i(t, s_i)$, where $t$ is calendar age in years and $s_i$ is sex (1: male, 0: female)), and another specifying the hazard rate (i.e., mortality rate in our applications) as a function of $t$, $s_i$, and the stochastic process $Y_i(t, s_i)$, denoted as $\mu\big(t, s_i, Y_i(t, s_i)\big)$:

$$\mathrm{d}Y_i(t, s_i) = \mathrm{a}(t, s_i)\big(Y_i(t, s_i) - \mathrm{f}_1(t, s_i)\big)\mathrm{d}t + \mathrm{b}(t, s_i)\mathrm{d}W_i(t), \text{ with initial condition } Y_i(t_0, s_i)$$

$$\mu\big(t, s_i, Y_i(t, s_i)\big) = \mu_0(t, s_i) + \mathrm{Q}(t, s_i)(Y_i(t, s_i) - \mathrm{f}_0(t, s_i))^2 I(Y_i(t, s_i) < \mathrm{f}_0(t, s_i))$$

Here, $W_i(.)$ denotes the Wiener process (Brownian motion), introducing stochasticity into the model; $I(.)$ is an indicator function; $b(.)$ is the diffusion (volatility) coefficient specifying the variability of the stochastic process; $f_1(.)$ represents the "equilibrium" EC trajectory (the long-term mean of $Y(.)$), reflecting the aging-related changes in ECs (for simplicity, we refer to this as "the estimated aging-related changes in ECs" in the text); $a(.)$ is the negative feedback coefficient regulating the dynamic behavior of $Y(.)$ (from a technical standpoint, negativity is required to ensure that the stochastic process does not deviate indefinitely from $f_1(.)$, and the absolute value of this coefficient determines the rate at which the process returns to the equilibrium level); $f_0(.)$ denotes the ECT that triggers the association of EC with mortality when ECs fall below it; $\mu_0(.)$ is the baseline hazard specifying mortality risk independent of EC; and $Q(.)$ is the non-negative scaling parameter in the quadratic hazard term that controls the width of the hazard function with respect to $Y(.)$. In essence, the modified SPM assumes that for ECs above the threshold $f_0(.)$, mortality risk equals the baseline level $\mu_0(.)$. If EC falls below $f_0(.)$, the mortality association is activated, and an additional term $\mathrm{Q}(t, s_i)(Y_i(t, s_i) - \mathrm{f}_0(t, s_i))^2$ appears in the hazard function.

We used parametric specifications of the SPM components similar to those employed in our recent applications to LLFS data [3]:

1) <u>Baseline hazard</u> $\mu_0(t, s_i)$: $\ln \mu_0(t, s_i) = \ln a_{\mu_0} + b_{\mu_0}(t - 70) + \beta_{\mu_0} s_i$, where $a_{\mu_0}$ is the baseline hazard rate for 70-year-old females (i.e., $s_i = 0$), $b_{\mu_0}$ represents the exponential growth of mortality with age, and $\beta_{\mu_0}$ is the parameter associated with sex in the baseline hazard.
2) <u>Scaling parameter in the quadratic hazard term</u> $\mathrm{Q}(t, s_i)$: Based on preliminary investigations, we used a sex-independent form because the available sample does not allow reliable estimation of sex-specific versions. We therefore specified $\mathrm{Q}(t, s_i) = a_{\mathrm{Q}} + b_{\mathrm{Q}}(t - 70)$, where $a_{\mathrm{Q}}$ corresponds to the baseline width of the parabola at age 70 (i.e., the shape of the EC-related mortality below the ECT for 70-year-olds), and $b_{\mathrm{Q}}$ models how this width changes with age (e.g., narrows if $b_{\mathrm{Q}} > 0$).
3) <u>Negative feedback coefficient</u> $\mathrm{a}(t, s_i)$: $a(t, s_i) = a_{\mathrm{Y}} + b_{\mathrm{Y}}(t - 70) + \beta_{\mathrm{Y}} s_i$, where $a_{\mathrm{Y}} < 0$, $b_{\mathrm{Y}} \geq 0$, $a_{\mathrm{Y}}$ represents the baseline value of this coefficient for 70-year-old females, $b_{\mathrm{Y}}$ models its rate of change with age, and $\beta_{\mathrm{Y}}$ quantifies the difference in baseline levels between males and females.
4) <u>Volatility coefficient</u> $\mathrm{b}(t, s_i)$: Following our earlier studies [2], we used an age-independent specification, $b(t, s_i) = \sigma_1 + \beta_{\mathrm{W}} s_i$, where $\sigma_1$ represents the volatility in females, and $\sigma_1 + \beta_{\mathrm{W}}$ is the corresponding coefficient in males.
5) <u>Equilibrium EC trajectory</u> $\mathrm{f}_1(t, s_i)$: Based on empirical observations indicating that the rate of decline of EC with age differs between females and males, we modeled the trajectories as $f_1(t, s_i) = a_{\mathrm{f}_1} + b_{\mathrm{f}_1}(t - 70) + \beta_{\mathrm{f}_1} s_i + \beta_{\mathrm{f}_1 s} s_i(t - 70)$. Here, $a_{\mathrm{f}_1}$ is the value at age 70 in

females; $b_{\mathrm{f}_1}$ describes the rate of age-related change in females; $\beta_{\mathrm{f}_1}$ determines the difference in at age 70 between males and females; and $\beta_{\mathrm{f}_1 s}$ quantifies the difference between males and females in the rate of age-related changes in EC.

6) <u>ECT</u> $\mathrm{f}_0(t, s_i)$: $f_0(t, s_i) = a_{\mathrm{f}_0} + b_{\mathrm{f}_0}(t - 70) + \beta_{\mathrm{f}_0} s_i$, where $a_{\mathrm{f}_0}$ is the ECT at age 70 in females, $b_{\mathrm{f}_0}$ defines its rate of change with age, and $\beta_{\mathrm{f}_0}$ reflects the difference in the ECT at age 70 between males and females.

We tested the following null hypotheses ($H_0$) using the likelihood ratio test, as in our prior work [3,4]: a) $H_0$: $\mathrm{Q}(t, s_i) = 0$ (i.e., there is no quadratic term in the hazard equation and, therefore, no EC-related mortality below the ECT); b) $H_0$: $b_{\mathrm{Q}} = 0$ (i.e., the width of the parabola describing EC-related mortality below the ECT does not change with age); c) $H_0$: $b_{\mathrm{f}_1} = 0$ and $b_{\mathrm{f}_1 s} = 0$ (i.e., no aging-related changes in ECs); d) $H_0$: $b_{\mathrm{f}_1 s} = 0$ (i.e., no difference in age-related EC changes between males and females); e) $H_0$: $b_{\mathrm{f}_0} = 0$ (i.e., the ECT does not change with age); f) $H_0$: $\beta_{\mu_0} = 0$, $\beta_{\mathrm{Y}} = 0$, $\beta_{\mathrm{W}} = 0$, $\beta_{\mathrm{f}_1} = 0$, $\beta_{\mathrm{f}_1 s} = 0$, and $\beta_{\mathrm{f}_0} = 0$ (i.e., all model components are the same in females and males); g) $H_0$: $\beta_{\mu_0} = 0$ (i.e., baseline mortality rates do not differ between females and males); h) $H_0$: $\beta_{\mathrm{Y}} = 0$ (i.e., the rate at which EC returns to equilibrium is the same in females and males); i) $H_0$: $\beta_{\mathrm{W}} = 0$ (i.e., EC variability is the same in both sexes); j) $H_0$: $\beta_{\mathrm{f}_1} = 0$ and $\beta_{\mathrm{f}_1 s} = 0$ (i.e., equilibrium EC trajectories are sex-independent); and k) $H_0$: $\beta_{\mathrm{f}_0} = 0$ (i.e., the ECT does not differ between females and males).

### Supplementary Tables

**Supplementary Table S1.** Detailed characteristics of the Long Life Family Study samples with one or more erythrocyte count (EC) measurements (EC1+) and two or more EC measurements (EC2+) used in the analyses

| Characteristics | EC1+ | EC2+ |
|---|---|---|
| Number of participants | 1,620 | 520 |
| Number (%) of deceased participants | 765 (47.2%) | 142 (27.3%) |
| Follow-up period (years) (mean ± SD [range]) | 10.3 ± 5.3 [0.00, 18.41] | 13.9 ± 2.9 [5.00, 18.41] |
| Age at first observation (years) (mean ± SD [range]) | 82.0 ± 9.2 [70, 110] | 80.8 ± 8.1 [70, 110] |
| Females (%) | 57.8% | 57.3% |
| US participants (%) | 66.6% | 56.0% |
| Smokers (smoked >100 cigarettes in lifetime) (%) | 39.9% | 39.0% |
| BMI at baseline (mean ± SD [range]), kg/$m^2$ | 26.8 ± 4.5 [15, 57] | 27.1 ± 3.9 [19, 42] |
| hsCRP (mean ± SD [range]), mg/L | 3.5 ± 8.8 [0.1, 147.8] | 2.9 ± 8.4 [0.1, 147.8] |
| CysC (mean ± SD [range]), mg/L | 1.1 ± 0.4 [0.5, 3.4] | 1.0 ± 0.3 [0.5, 2.2] |
| IL-6 (mean ± SD [range]), pg/mL | 2.5 ± 6.7 [0.1, 116.4] | 1.9 ± 5.3 [0.2, 92.2] |
| sTfR (mean ± SD [range]), mg/L | 3.0 ± 1.1 [1.2, 18.4] | 2.8 ± 0.7 [1.3, 7.5] |
| EC (mean ± SD [range]), $10^{12}$/L, ages 70 years and older | 4.48 ± 0.51 [2.20, 8.00] | 4.51 ± 0.47 [2.50, 6.40] |
| EC (mean ± SD [range]), $10^{12}$/L, ages 70-79 years | 4.64 ± 0.46 [2.20, 8.00] | 4.63 ± 0.42 [3.20, 6.40] |
| EC (mean ± SD [range]), $10^{12}$/L, ages 80-89 years | 4.45 ± 0.45 [2.50, 5.90] | 4.47 ± 0.46 [2.90, 5.90] |
| EC (mean ± SD [range]), $10^{12}$/L, ages 90-99 years | 4.26 ± 0.52 [2.50, 6.30] | 4.24 ± 0.50 [2.50, 5.60] |
| EC (mean ± SD [range]), $10^{12}$/L, ages 100 years and older | 4.02 ± 0.58 [2.30, 5.50] | 3.99 ± 0.46 [3.00, 4.80] |
| Annual rate of change in EC (mean ± SD [range]), $10^{12}$/L per year, ages 70 years and older | | -0.020 ± 0.056 [-0.271, 0.186] |
| Annual rate of change in EC (mean ± SD [range]), $10^{12}$/L per year, ages 70-79 years | | -0.011 ± 0.052 [-0.171, 0.186] |
| Annual rate of change in EC (mean ± SD [range]), $10^{12}$/L per year, ages 80-89 years | | -0.024 ± 0.069 [-0.271, 0.100] |
| Annual rate of change in EC (mean ± SD [range]), $10^{12}$/L per year, ages 90 years and older | | -0.031 ± 0.076 [-0.240, 0.150] |

BMI – body mass index, CysC – cystatin C, EC – erythrocyte count, hsCRP – high sensitivity C-reactive protein, IL-6 – interleukin-6, sTfR – transferrin receptor, SD – standard deviation. 2) Annual rate of change in EC: For participants with two visits, the rate is computed as $\Delta EC = (EC_{visit2} - EC_{visit1})/(Age_{visit2} - Age_{visit1})$; for participants with three visits, values of $\Delta EC$ are computed separately for the intervals between visit 2 and visit 1, and between visit 3 and visit 2. 3) Sample sizes: Actual sample sizes vary across models because of missing values for different variables. The following numbers of missing values were observed: hsCRP – 118, CysC – 25, IL-6 – 47, sTfR – 174).

**Supplementary Table S2.** Results of the joint model adjusted for basic demographic characteristics, applied to data on mortality and erythrocyte counts in LLFS participants aged 70 years or older

#### *A. Longitudinal submodel*

| Variable | beta | SE | CI | p |
|---|---|---|---|---|
| Intercept | 5.882 | 0.094 | [5.691, 6.071] | $< 1 \times 10^{-300}$ |
| age1 | -0.018 | 0.001 | [-0.021, -0.016] | $1.2 \times 10^{-58}$ |
| sex | 0.192 | 0.023 | [0.144, 0.238] | $8.1 \times 10^{-17}$ |
| time | -0.021 | 0.002 | [-0.025, -0.016] | $8.2 \times 10^{-18}$ |

#### *B. Survival submodel*

| Variable | beta | SE | HR | CI | p |
|---|---|---|---|---|---|
| $EC_{ind}$ | -0.326 | 0.108 | 1.386 | [1.177, 1.691] | 0.003 |
| age1 | 0.135 | 0.003 | 1.145 | [1.134, 1.154] | $< 1 \times 10^{-300}$ |
| sex | 0.348 | 0.074 | 1.416 | [1.241, 1.594] | $2.8 \times 10^{-6}$ |

**A.** Estimates from the longitudinal submodel, which models individual erythrocyte count (EC) trajectories. Columns: **beta** – estimate of the regression coefficient for the variable listed in **Variable**; **SE** – standard error of beta; **CI** – 95% bootstrap confidence interval for the beta; **p** – p-value for the null hypothesis beta = 0 for respective variables: intercept, age at first EC observation of (age1), sex (1 – males, 0 – females), and time – time since the first EC measurement. **B.** Estimates from the survival submodel, which relates characteristics of individual EC trajectories to mortality risk. Columns: **beta** – estimate of the regression coefficient for the variable listed in **Variable**; **SE** – standard error of beta; **HR** – hazard ratio for a one-unit *increase* in the respective variable (except for $EC_{ind}$, the subject-specific deviation from expected EC; for $EC_{ind}$, HR corresponds to a one-unit *decrease*); **CI** – 95% bootstrap confidence interval for the HR; and **p** – p-value for the null hypothesis beta = 0 for the respective variable. Confidence intervals were obtained using the familial bootstrap, whereas p-values are from the fitted *joineRML* model output; therefore, exact agreement between the two inferential summaries is not expected.

**Supplementary Table S3.** Results of the joint model adjusted for basic demographic characteristics and age-time and sex-time interaction terms, applied to data on mortality and erythrocyte counts in LLFS participants aged 70 years and older

***A. Longitudinal submodel***

| Variable | beta | SE | CI | p |
|---|---|---|---|---|
| Intercept | 5.519 | 0.122 | [5.300, 5.748] | $< 1 \times 10^{-300}$ |
| age1 | -0.014 | 0.001 | [-0.017, -0.011] | $7.5 \times 10^{-21}$ |
| sex | 1.148 | 0.189 | [0.654, 1.509] | $1.2 \times 10^{-9}$ |
| time | -0.016 | 0.003 | [-0.022, -0.010] | $1.2 \times 10^{-6}$ |
| age1*sex | -0.012 | 0.002 | [-0.016, -0.005] | $2.3 \times 10^{-7}$ |
| time*sex | -0.011 | 0.005 | [-0.018, -0.003] | 0.014 |

***B. Survival submodel***

| Variable | beta | SE | HR | CI | p |
|---|---|---|---|---|---|
| $EC_{ind}$ | -0.365 | 0.113 | 1.441 | [1.201, 1.755] | 0.001 |
| age1 | 0.136 | 0.003 | 1.145 | [1.134, 1.155] | $< 1 \times 10^{-300}$ |
| sex | 0.375 | 0.072 | 1.455 | [1.271, 1.648] | $2.1 \times 10^{-7}$ |

**A.** Estimates from the longitudinal submodel, which models individual erythrocyte count (EC) trajectories. Columns: **beta** – estimate of the regression coefficient for the variable listed in **Variable**; **SE** – standard error of beta; **CI** – 95% bootstrap confidence interval for the beta; **p** – p-value for null hypothesis beta = 0 for respective variables: intercept, age at first EC measurement (age1), sex (1 – males, 0 – females), time – time since the first EC measurement, age1*sex – age1-by-sex interaction, and time*sex – time-by-sex interaction. **B.** Estimates from the survival submodel, which relates characteristics of individual EC trajectories with mortality risk. Columns: **beta** – estimate of the regression coefficient for the variable listed in **Variable**; **SE** – standard error of beta; **HR** – hazard ratio for a one-unit *increase* in the respective variable, except for $EC_{ind}$ (the subject-specific deviation from expected EC, see **Methods**), for which the HR corresponds to a one-unit *decrease*; **CI** – 95% bootstrap confidence interval for the HR; and **p** – p-value for the null hypothesis beta = 0 for the respective variable. Confidence intervals were obtained using the familial bootstrap, whereas p-values are from the fitted *joineRML* model output; therefore, exact agreement between the two inferential summaries is not expected.

**Supplementary Table S4.** Results of the Cox proportional hazards model adjusted for additional covariates, applied to data on mortality and the first observation of erythrocyte counts in LLFS participants aged 70 years and older

| **Variable** | **beta** | **SE** | **HR** | **CI** | **p** |
|---|---|---|---|---|---|
| EC | -0.312 | 0.091 | 1.366 | [1.130, 1.678] | $7.0 \times 10^{-4}$ |
| sex | 0.309 | 0.082 | 1.362 | [1.227, 1.614] | $1.1 \times 10^{-4}$ |
| IsDK | 0.314 | 0.095 | 1.368 | [1.114, 1.628] | $7.8 \times 10^{-4}$ |
| BMIbin | -0.243 | 0.083 | 0.784 | [0.668, 0.905] | 0.002 |
| Cysc | 0.412 | 0.065 | 1.510 | [1.352, 1.771] | $1.4 \times 10^{-10}$ |
| hsCRP | 0.077 | 0.043 | 1.080 | [0.981, 1.177] | 0.104 |
| Il6bin | 0.250 | 0.122 | 1.284 | [1.019, 1.579] | 0.050 |
| Smoke100 | 0.088 | 0.083 | 1.092 | [0.940, 1.286] | 0.301 |
| sTfR | 0.160 | 0.042 | 1.174 | [1.083, 1.274] | $2.0 \times 10^{-4}$ |

Variables: EC – erythrocyte count; sex (1 – males, 0 – females); IsDK – country (1 – Denmark, 0 – USA); BMIbin – body mass index (1 – above median, 0 – below median); CysC – cystatin C; hsCRP – high sensitivity C-reactive protein; IL6bin – interleukin-6 (1 – above median, 0 – below median); Smoke100 – smoked >100 cigarettes in lifetime; sTfR – transferrin receptor. Columns: **beta** – estimate of the regression coefficient for the variable listed in **Variable**; **SE** – standard error of beta; **HR** – hazard ratio for a one-unit *increase* in the respective variable (except for EC, where HR corresponds to a one-unit *decrease*); **CI** – 95% bootstrap confidence interval for the HR; **p** – p-value for the null hypothesis beta = 0 for the respective variable. Note that age is used as a strata variable (with strata: 70-74, 75-79, 80-84, 85-89, and 90+ years; see **Methods**). Therefore, age variable is not shown in the table.

**Supplementary Table S5.** Results of the Cox proportional hazards model adjusted for basic demographic characteristics and additional covariates, applied to data on mortality and the first observation of erythrocyte counts in LLFS participants from different age groups

| Age Group | beta | SE | HR | CI | p | N |
|---|---|---|---|---|---|---|
| 70+ | -0.312 | 0.091 | 1.366 | [1.130, 1.678] | $7.0 \times 10^{-4}$ | 1403 |
| 75+ | -0.299 | 0.092 | 1.349 | [1.107, 1.630] | 0.001 | 1107 |
| 80+ | -0.320 | 0.096 | 1.377 | [1.144, 1.683] | $6.2 \times 10^{-4}$ | 838 |
| 85+ | -0.351 | 0.101 | 1.420 | [1.169, 1.745] | $4.1 \times 10^{-4}$ | 633 |
| 90+ | -0.372 | 0.109 | 1.450 | [1.132, 1.767] | $2.9 \times 10^{-4}$ | 449 |

Estimates for the erythrocyte count (EC) variable in the Cox model, analogous to that shown in Supplementary Table S4, applied to LLFS participants in different age groups (**Age Group**). **N** denotes the sample size (note that the sample size in the 70 years or older group differs from that in the basic Cox model reported in the main text because individuals with missing values for any of the additional variables were removed). All other columns are defined as in Supplementary Table S4.

**Supplementary Table S6.** Results of the joint model adjusted for basic demographic characteristics and additional covariates, applied to data on mortality and erythrocyte counts in LLFS participants aged 70 years and older

***A. Longitudinal submodel***

| Variable | beta | SE | CI | p |
|---|---|---|---|---|
| Intercept | 5.311 | 0.166 | [4.992, 5.637] | $1.6 \times 10^{-224}$ |
| age1 | -0.011 | 0.002 | [-0.015, -0.007] | $8.2 \times 10^{-8}$ |
| sex | 1.291 | 0.205 | [0.775, 1.662] | $3.3 \times 10^{-10}$ |
| age1*sex | -0.014 | 0.002 | [-0.018, -0.007] | $4.2 \times 10^{-8}$ |
| time | -0.016 | 0.003 | [-0.023, -0.011] | $1.7 \times 10^{-6}$ |
| time*sex | -0.010 | 0.005 | [-0.018, -0.001] | 0.032 |
| IsDK | 0.050 | 0.028 | [0.007, 0.117] | 0.075 |
| BMIbin | 0.034 | 0.026 | [-0.030, 0.085] | 0.180 |
| hsCRP | 0.002 | 0.013 | [-0.023, 0.027] | 0.887 |
| Cysc | -0.021 | 0.018 | [-0.055, 0.009] | 0.230 |
| Il6bin | -0.038 | 0.033 | [-0.083, 0.016] | 0.240 |
| Smoke100 | -0.025 | 0.026 | [-0.065, 0.011] | 0.346 |
| sTfR | 0.061 | 0.011 | [0.035, 0.081] | $8.1 \times 10^{-8}$ |

***B. Survival submodel***

| Variable | beta | SE | HR | CI | p |
|---|---|---|---|---|---|
| $EC_{ind}$ | -0.333 | 0.128 | 1.395 | [1.157, 1.774] | 0.009 |
| age1 | 0.115 | 0.005 | 1.122 | [1.109, 1.136] | $2.0 \times 10^{-102}$ |
| sex | 0.341 | 0.085 | 1.407 | [1.255, 1.642] | $5.8 \times 10^{-5}$ |
| IsDK | 0.308 | 0.099 | 1.360 | [1.126, 1.606] | 0.002 |
| BMIbin | -0.248 | 0.084 | 0.780 | [0.668, 0.900] | 0.003 |
| hsCRP | 0.062 | 0.043 | 1.063 | [0.971, 1.154] | 0.154 |
| Cysc | 0.367 | 0.064 | 1.443 | [1.275, 1.707] | $9.4 \times 10^{-9}$ |
| Il6bin | 0.206 | 0.129 | 1.229 | [1.009, 1.504] | 0.111 |
| Smoke100 | 0.133 | 0.084 | 1.142 | [0.998, 1.351] | 0.115 |
| sTfR | 0.155 | 0.040 | 1.167 | [1.079, 1.259] | $9.8 \times 10^{-5}$ |

Estimates from the longitudinal submodel, which models individual erythrocyte count (EC) trajectories. Columns: **beta** – estimate of the regression coefficient for the variables listed in **Variable**; **SE** – standard error of beta; **CI** – 95% bootstrap confidence interval for the beta; **p** – p-value for the null hypothesis beta = 0 for respective variables: intercept; age1 – age at first EC measurement; sex (1 – males, 0 – females); age1*sex – age1-by-sex interaction; time – time since the first EC measurement; time*sex – time-by-sex interaction; IsDK – country (1 – Denmark, 0 – USA); BMIbin – body mass index (1 – above median, 0 – below median); CysC – cystatin C; hsCRP – high sensitivity C-reactive protein; IL6bin – interleukin-6 (1 – above median, 0 – below median); Smoke100 – smoked >100 cigarettes in lifetime; sTfR – transferrin receptor. **B.** Estimates from the survival submodel, which relates characteristics of individual EC trajectories to mortality risk. Columns: **beta** – estimate of the regression coefficient for the variable listed in **Variable**; **SE** – standard error of beta; **HR** – hazard ratio for a one-unit *increase* in the respective variable, except for $EC_{ind}$ (the subject-specific deviation from the expected EC; see **Methods**), for which the HR corresponds to a one-unit *decrease*; **CI** – 95% bootstrap confidence interval for the HR; **p** – p-value for the null hypothesis beta = 0 for the respective variable. Note that the time since the first EC measurement is used as the model's time variable. Therefore, the time variable does not appear in the survival submodel. Confidence intervals were obtained using the familial bootstrap, whereas p-values are from the fitted *joineRML* model output; therefore, exact agreement between the two inferential summaries is not expected.
